# Tandem repeat variation within and between species reveals signatures of selection in humans and chimpanzees

**DOI:** 10.64898/2026.01.20.700717

**Authors:** Carolina L. Adam, Joana L. Rocha, Peter H. Sudmant, Rori V. Rohlfs

## Abstract

Tandem repeats (TRs) are highly mutable DNA elements that influence gene regulation^1^, protein structure^2^, and disease^3^. Until recently, their repetitive nature has hindered accurate TR sequencing and genotyping, resulting in sparse comparative data across species. In addition, we lack population-aware approaches to analyze TR conservation, divergence, and mutational dynamics. Here, leveraging telomere-to-telomere primate genomes and long-read data from 46 humans and 23 chimpanzees, we constructed a catalog of homologous TR loci, and developed an analytical framework to jointly analyze TR variation within- and between-species. Across primates, TR diversity and conservation vary strongly with genomic context, with coding and 5’ UTR TRs exhibiting reduced polymorphism and constraint across species, consistent with stabilizing selection. Yet, while TRs are depleted in coding sequence, they are enriched in 5’ UTRs, suggesting functional roles that outweighs mutational risks. TR heterozygosity varies across motif lengths and is concordant with both evolutionary and trio-based mutation rate estimates^4^. Introducing an HKA-like approach to control for locus-specific mutation rates, we identified TRs with signatures of directional and balancing selection. These candidates are significantly enriched in genes involved in nervous system development and synaptic function, highlighting TRs as potential contributors to neural evolution. Further, TR divergence correlates with gene expression divergence, particularly for promoter-related TRs and expression in organoids related to neurodevelopment, implicating a subset of regulatory TRs as candidates for adaptive expression evolution. Finally, trait-associated TRs display longer alleles and higher diversity in humans compared to chimpanzees, consistent with lineage-specific runaway mutations and/or directional selection^5^. Together, our results establish a comparative framework for TR evolutionary analyses, revealing how mutational processes and selection jointly shape repeat variation, and supporting their role as both conserved functional elements and as drivers of evolutionary innovation.

## Main

Tandem repeats (TRs) are ubiquitous across metazoans and account for nearly 8% of the human genome^6^. Their repetitive structure makes them prone to replication slippage, leading to mutation rates that are orders of magnitude higher than single-nucleotide variants (SNVs)^7^. On a per-locus basis, TR variants are more likely than SNVs to impact functional traits^8^, and human genetics has repeatedly demonstrated their impact on development and disease, from neurological disorders to cancer^9–11^. These properties suggest that TRs may act as fuel for rapid adaptation^12,13^.

Despite multiple lines of evidence supporting the role of TRs in adaptation^13–16^, technical limitations in sequencing and genotyping long repetitive DNA have hindered systematic TR analyses across species. Now, advances in long-read sequencing and telomere-to-telomere (T2T) assemblies have resolved previously inaccessible repeat regions^6^, fundamentally reshaping our understanding of genome architecture and structural variation^17^. As a result, recent comparative studies have uncovered lineage-specific TRs associated with evolutionary divergence among primates, particularly in neuron-specific regulatory mechanisms^18–20^. Characterizing TR variation between species can provide insight into how TRs impact molecular processes by revealing conserved sites under stabilizing selection, as well as sites under directional selection that underlie adaptive lineage-specific traits. Still, we lack a framework that enables comparisons of long-read TR genotypes both between and within species, an approach that has proven transformative in SNV-based evolutionary studies^21,22^.

Here, we leveraged T2T reference genomes to identify millions of homologous TR loci between humans and non-human primates (NHPs), revealing broad patterns of TR constraint over evolutionary time. We integrated long-read, assembly-level data from 46 humans, and 23 newly sequenced chimpanzees, described in a companion paper^23^, providing a unique resource for joint analysis of within- and between-species variation to investigate the mutational and selective processes shaping TR evolution. Further, we propose an analytical HKA-like framework for comparative TR analyses, enabling the identification of locus-specific deviations in divergence-diversity ratios and candidate loci under distinct selective regimes. Finally, we examine the contribution of TR variation to expression divergence and trait variation, highlighting how extreme patterns of divergence and diversity may help prioritize putatively functional loci.

## Results

### Species-specific TR catalogs, homology to humans, and genomic distribution

We developed the TRACK pipeline to create species-specific TR catalogs and identify homology between human and each NHP reference genome. TRs are subdivided based on their motif length, where short tandem repeats (STRs) exhibit motifs ≤ 6 bp, and variable number tandem repeats (VNTRs) display motifs ≥ 7bp. Their distributions across motif lengths and catalog sizes were comparable across species (Fig. 1a-c). Among STRs, abundance across motif lengths was highly concordant with other studies (Fig. 1c)^24–26^. Discordances, such as hexamer abundance^24,25^ where we observed a mild depletion, likely stem from differences in TR identification and filtering. For instance, while some studies consider only TRs with high or complete sequence constancy^24–26^, our analysis allowed a lower threshold (>60%), capturing more variable repeats. Motifs larger than 20 bp were considerably less common and have not been systematically explored in prior studies. When queried in the Dfam database, several common long motifs matched known repetitive elements, including Alu elements, and VNTRs embedded within composite retrotransposons (Fig. 1a). Young Alu elements have been reported to insert adjacent to older Alus, potentially through reuse of nearby LINE-1 cleavage sites^27^. The GC content of TR motifs also varied with motif lengths, but showed broadly consistent patterns across species (Supplementary Fig. 2).

**Fig. 1.**
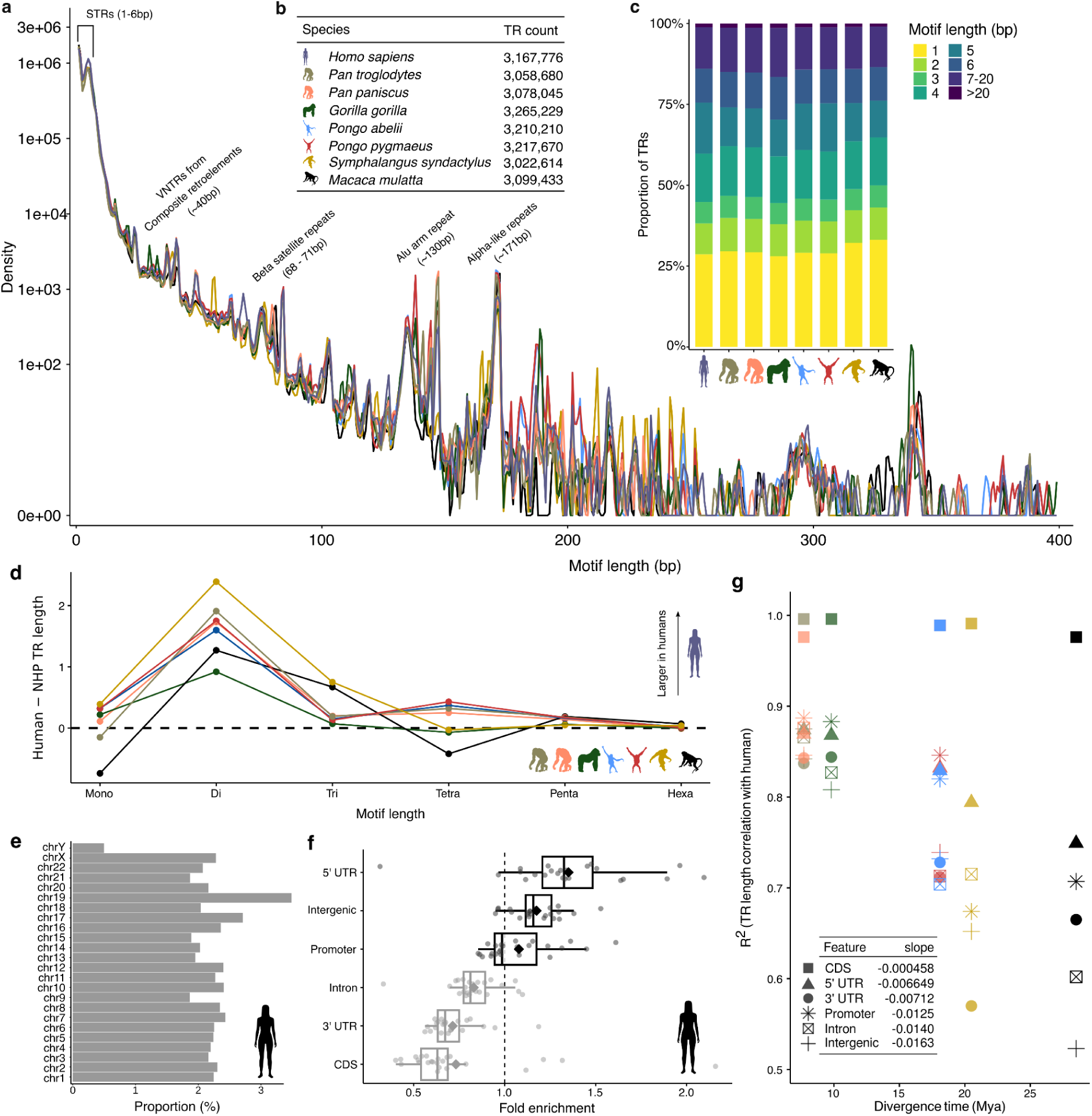
Tandem repeats across T2T primate genomes. **a**, Distributions of TR motif lengths across primate genomes. Labeled peaks indicate TRs with motif lengths >20 bp that show strong matches to entries in the Dfam database. **b**, TR count per species. **c**, Proportional distributions of STR motif lengths across primates. **d**, Difference in total STR lengths between human and each NHP species across motif lengths. **e**, Proportion of each human chromosome contained in TRs. **f**, Fold enrichment of total TR length (bp) overlapping annotated genomic features per chromosome. Diamonds indicate mean enrichment across the genome. **g**, Divergence time (in millions of years) and the coefficient of determination (R²) for reference TR length comparisons between humans and each non-human primate species. The inset table shows the slope of the linear regression line for each genomic feature.

Total STR length varies significantly across motif sizes and between species (Fig. 1d; Supplementary Table S1). The largest differences were observed for dimers, which were consistently longer in humans than in all NHP species, consistent with earlier human-chimpanzee results^28^. The persistence of this pattern across a broader primate phylogeny is consistent with a shift on the human lineage for an increased rate of dinucleotide expansions.

STRs were generally uniformly distributed along chromosome lengths (Extended Data Fig. 1). In humans, chromosomes 19 and 17 showed the highest relative TR densities (Fig. 1e), consistent with earlier studies^29,30^. Chromosome Y displayed the lowest density of non-CenSat TRs after filtering (see Methods), reflecting the exclusion of its extensive satellite-rich regions^31^. In contrast to STRs, human VNTRs are concentrated in subtelomeric regions (Extended Data Fig. 1a), in line with previous observations that subtelomeric structural variation in humans is largely composed of VNTRs with motifs >15 bp^32^. In several NHP, however, telomeric and subtelomeric regions are depleted of TRs after filtering (Extended Data Fig. 1b-h). This reflects the presence of lineage-specific satellite-like repeat structures at chromosome ends, including the StSats arrays described in non-human great apes^33^ and the large SiaRep repeats found in siamang gibbons^34^, which were masked during filtering. The density of TRs across homologous genomic regions shows decreasing correlation with increasing time since the most recent common ancestor (TMRCA), indicating progressive divergence of repeat landscapes over evolutionary time (Extended Data Fig. 2).

In humans, TRs are depleted in coding (CDS) regions, 3’ untranslated regions (UTRs), and introns, while they are enriched in 5’ UTRs, intergenic regions, and promoters (Fig. 1f). When stratified by motif length, distinct distributions emerged: (Supplementary Fig. 3, Supplementary Table S2), in agreement with a large STR population panel^12^. Monomers and dimers were strongly depleted in coding regions (0.01 and 0.05-fold, respectively), while repeats with motif lengths in multiples of three were overrepresented (Supplementary Fig. 3, Supplementary Table S2), reflecting the coding-frame constraint^25,30^. A similar trend was observed in the 5’ UTRs. The GC content of TR motifs also varied across genomic features, being enriched in coding and 5’ UTRs (Supplementary Fig. 4).

The varying abundance of homologous TRs between human and NHPs reflects phylogenetic distances, with chimpanzees retaining the largest overlap (Supplementary Fig. 5). TR length comparison between species reveals conservation following the phylogeny, with correlations weakening as TMRCA increases, with higher conservation for TRs within CDS (Fig. 1g, Extended Data Fig. 3). In coding regions, the maintenance of TR length has been shown to be crucial in several cases to preserve functional protein structure^35,36^, leading to deep conservation of coding TRs across mammals^37^. We also observe notable length conservation for TRs in 5’ UTRs (Fig. 1g, Extended Data Fig. 3). In 5’ UTRs, extreme conservation has been linked to RNA-mediated translational control in essential developmental genes^38^. In accordance, genes with TRs in their 5’UTRs are gene ontology (GO) enriched with biological processes spanning broad terms such as developmental growth, to more specialized terms such as axonogenesis, with overall enrichment for core neurodevelopmental processes (Supplementary Fig. 6). Additionally, variants in UTRs are known to disrupt transcription initiation and, in some cases, contribute to disease^39^.

### TR genotypes, comparative length variation, and mutation process

After quality control and filtering (see Methods), we identified 1,905,929 homologous TRs between humans and chimpanzees. Of these, 1,033,666 (54.2%) are invariant, i.e., monoallelic across samples, and 872,237 (45.7%) exhibit at least two different alleles in either species, here denoted TR variants, or TRVs. Invariant TRs display longer motifs and have an overrepresentation of coding and 5’ UTR repeats (Supplementary Fig. 7).

Homologous TRs are generally short (mean allele length ≤ 22 bp for 62.1% of TRs) and highly correlated between species (Fig. 2a-c; Supplementary Fig. 8). In coding and UTR regions, concordance is exceptionally high and mirrors previous assembly-level comparisons of orthologous human-chimpanzee STRs^40^, suggesting stronger stabilizing selection on functional TR variation. Highly divergent TRs also tend to exhibit greater allelic diversity within the species harboring the longer alleles (Fig. 2d; Supplementary Fig. 9). This pattern is consistent with longer alleles being subject to higher mutation rates and the potential for lineage-specific runaway repeat expansions^5,41^.

**Fig. 2.**
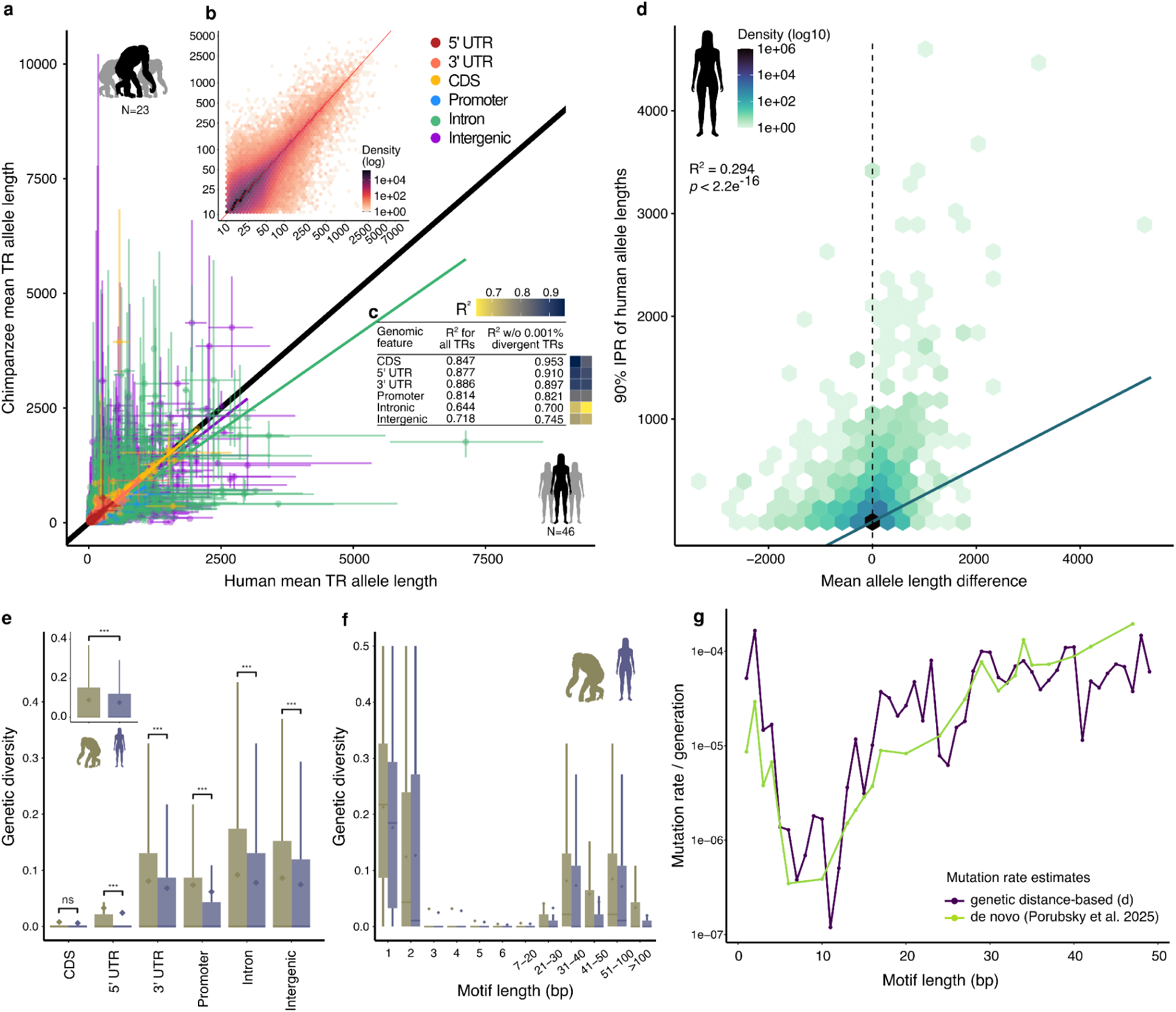
Comparative analyses of homologous tandem repeats between humans and chimpanzees. **a**, Mean TR allele lengths between humans (x-axis) and chimpanzees (y-axis) for TRs shared between species, including iTRs and TRVs. Each point represents a single TR locus, color-coded by its genomic annotation in the CHM13 genome. Whiskers represent the 5th-95th quantiles of allele lengths in humans (horizontal bars) and chimpanzees (vertical bars). **b**, Heatmap of the same TR loci showing the joint distribution of human and chimpanzee mean allele lengths. Color intensity reflects the density of TRs within each bin (50 bins). **c**, Correlation between human and chimpanzee TR length grouped by genomic feature. **d**, Heatmap of mean allele length divergence between chimpanzees and humans (x-axis) and the 90% interpercentile range of human allele lengths per locus. **e,f**, Expected heterozygosity across different (**e**) genomic features and (**f**) across motif lengths. \**p* ≤ 0.05, \*\**p* ≤ 0.01 and \*\*\**p* ≤ 0.001 denote statistically significant differences given by Wilcoxon rank-sum tests. **g**, Estimates of *de novo* (light green) and genetic distance-based (dark purple) mutation rates averaged across motif lengths.

Overall, chimpanzees show higher TR heterozygosity (Fig. 2e), reflecting lower human genetic diversity found across diverse markers^22,42,43^. Consistent with other studies^12,44^, TR genetic diversity is reduced in 5’ UTRs and coding regions in both species, suggesting that strong purifying selection limits functional TR variation across lineages. Heterozygosity also varied across motif lengths, highest for monomers and dimers, decreasing with motif lengths up to 20 bp, and increasing for longer motifs (Fig. 2f). This pattern holds for TRs across genomic features, except for coding and, to some extent, 5’ UTR regions, which have lower heterozygosity even for monomers and dimers (Supplementary Fig. 10). Thus, even when harboring short motifs known to have elevated mutation rates due to frequent slippage^7^, variation in coding, and to a lesser degree 5’ UTR, TRs are likely constrained by strong purifying selection. In 5’ UTRs, TRs exhibit remarkably low heterozygosity, reflecting the abundance of 3-6bp motifs, which display lower genetic diversity compared to monomers or dimers (Supplementary Figs. 3, 10). Nonetheless, TRs are overrepresented in 5’ UTRs compared to the genome-wide distribution (Fig. 2c; Odds ratio = 1.37, *p* < 2.2 × 10^−16^), and display even stronger enrichment in hyperconserved 5’ UTRs^38^ (Odds ratio = 3.54, *p* < 2.2 × 10^−16^), suggesting that they serve functional roles that offset the risk of mutations.

Comparative TR variation data also facilitates model-based estimates of locus-specific mutation rates. We estimated per locus mutation rates under the stepwise mutation model (SMM) and selective neutrality^45^ based on observed genetic distances between species. When stratified by TR motif length, our estimates are highly concordant with *de novo* mutation rates observed in human pedigree data^4^ (Fig. 2g). This concordance is particularly remarkable since SMM-based estimates overestimate mutation rates for loci which undergo multi-step mutations, and for the *de novo* observations we consider only mutation occurrence, not magnitude of change^46,47^. For short motifs where we have the most data, our SMM-based mutation rate estimates are higher than *de novo* observations. This may reflect frequent multistep mutations, as have been observed particularly for monomers and dimers^46,48^, or a TR mutation rate slow-down on the human lineage. Both mutation rate estimates generally decrease with motif length until about 20bp, when they increase. This pattern suggests a shift in the dominant mutational process, potentially from replication slippage in short motifs to recombination or gene conversion in longer motifs^49^. However, when considering only coding and regulatory TRs, model-based estimates drop below *de novo* mutation estimates, with concordance decreasing by an order of magnitude (Extended Data Fig. 4; MSE = 2.5 × 10^−8^ for intergenic TRs versus 1.4 × 10^−7^ for CDS TRs). This likely reflects violation of the assumption of selective neutrality at TRs subject to stronger stabilizing selection over evolutionary time. Further, we observe concordance between heterozygosity and mutation rate estimates over motif lengths, underscoring the role of mutational dynamics in shaping TR variation.

### TRs with exceptional divergence or diversity

Since TR mutation rate varies so widely across loci, allele length divergence between species of any given TR can arise from either species-specific selection or high mutation rates. We sought to account for this using an HKA-like approach by computing a species-specific divergence-diversity ratio (*D*) for each TR (See Methods), defined as the variance in TR length between species divided by the variance within the focal species. This approach controls for per-locus mutation rate variation by identifying loci with extreme divergence between species relative to within-species diversity. *D* values were broadly concordant between species (Fig. 3a; Pearson’s r = 0.15, *p* < 2.2 × 10^−16^; Spearman’s ⍴ = 0.84; *p* < 2.2 × 10^−16^), consistent with a general pattern of similar TRV mutational dynamics and selective pressures between humans and chimpanzees. To explore outlier TRVs as candidates for selection, we considered the 1,000 most extreme genic TRVs in each of three groups: 1) those with high *D* in both species, reflecting elevated divergence between species relative to within-species variation, consistent with species-specific directional selection; 2) those with low *D* in both species, characterized by relatively reduced divergence compared to within-species diversity, which may reflect balancing selection or strong stabilizing selection with a high mutation rate; and 3) those with asymmetric *D* patterns, where between-species divergence is coupled with higher diversity in either chimpanzees (chimp-biased *D*) or humans (human-biased *D*), which may be a product of directional selection^50,51^ or species-specific runaway mutations^36,52^ (Supplementary Table S3). Across groups, highly divergent genic TRVs were longer and with larger motifs (Wilcoxon rank-sum test, FDR-adjusted *p* < 0.0001; Supplementary Fig. 11a-d). They also displayed greater GC content when compared to a set of background TRVs with the same TR length and motif length distribution (Wilcoxon rank-sum test, FDR-adjusted *p* < 0.0001; Supplementary Fig. 11c) and were highly enriched in coding regions compared to the distribution of all genic TRVs (Odds ratio = 6.6, *p* < 2.2 × 10^−16^; Supplementary Fig. 11e).

**Fig. 3.**
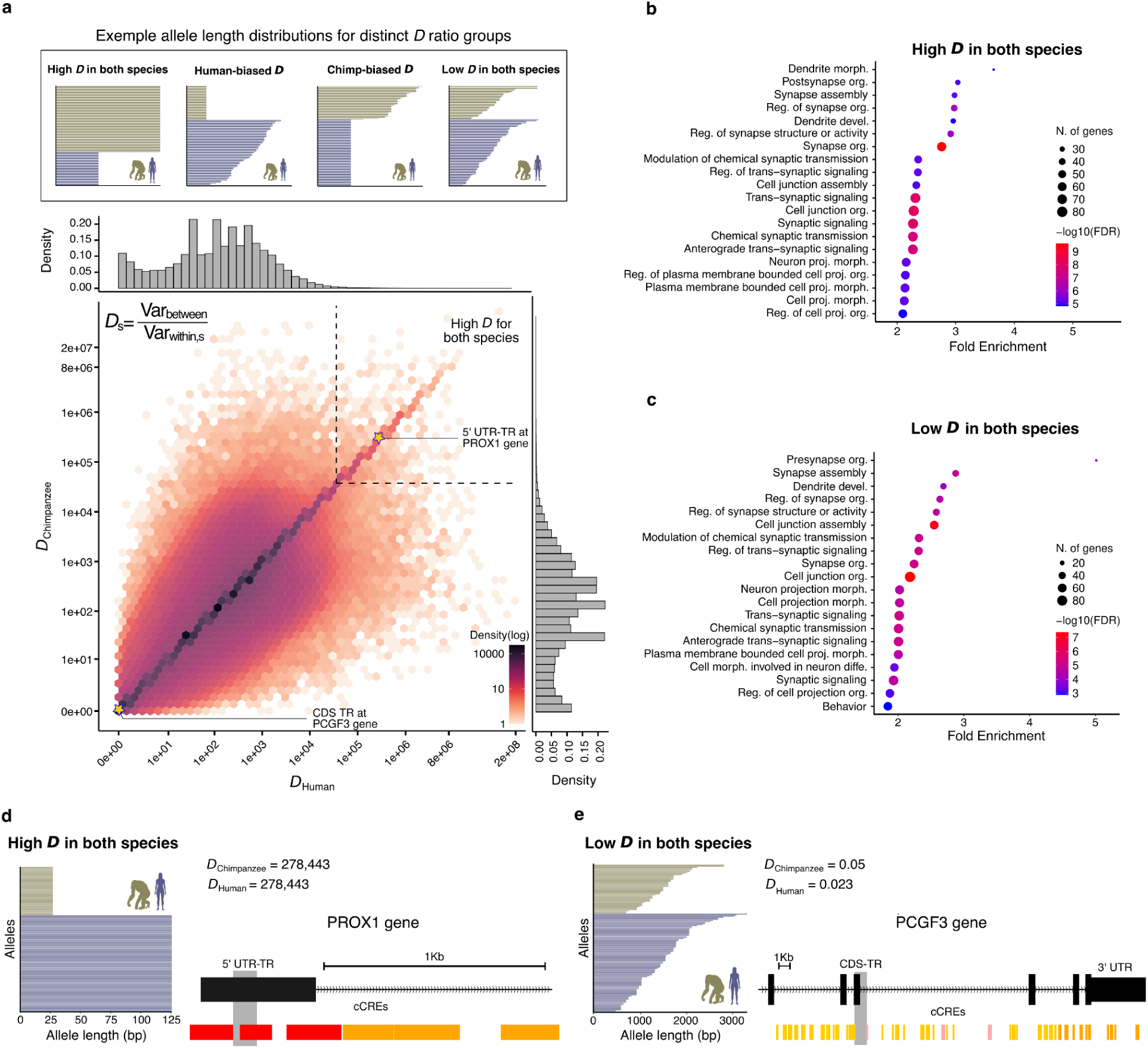
Divergence–diversity landscapes of TRs in humans and chimpanzees. **a**, Heatmap showing the joint distribution of TR Divergence-Diversity Ratios (*D*) in humans and chimpanzees. Marginal histograms display the distribution of *D* for each species. Black dashed line shows the boundaries of high *D* in both species. Stars indicate example TRs belonging to extreme ratio categories in both species. **b,c**, Gene Ontology enrichment analysis for biological processes terms associated with genes intersecting the top 1,000 genic TRs with (**b**) high *D* in both species and low *D* in both species. The set of all TR-containing genes was used as background. **d,e**, Genomic location and allele length distribution for two example TRs classified as (**d**) high *D* in both species and (**e**) low *D* in both species. Grey vertical bars represent TR location. Candidate cis-regulatory elements (cCREs) indicate promoter-like signature (red), proximal (orange), and distal (yellow) enhancer-like signature, and DNase-H3K4me3 elements (pink).

To investigate the biological processes associated with TRVs with extreme divergence or diversity, we performed GO enrichment analysis on the genes intersecting TRVs from each ratio group. While all TR-containing genes are modestly enriched for broad developmental processes (∼1.2-fold; Supplementary Fig. 12), genes containing both high and low *D* TRs showed >2-fold enrichment for terms underlying nervous system development, synaptic organization and cell signaling (Fig. 3b and c), despite their contrasting evolutionary patterns. For comparison, random sets of TR-containing genes showed no GO enrichment (see Supplementary Methods).

Of the 32 genes containing coding TRVs in the high *D* group, 13 are zinc-fingers (ZNF) genes, a functionally diverse family involved in processes such as transcriptional regulation and DNA repair^53^. Several ZNFs are known to undergo rapid lineage-specific divergence and positive selection on DNA-binding domains between humans and chimpanzees^54,55^. The strongest divergence signals from genic TRVs occur in introns of genes associated with neural and sensory systems, epithelial integrity, and signal transduction (Supplementary Table S4). One compelling candidate overlaps the 5’ UTR of PROX1 (Fig. 3d), a homeobox transcription factor essential for embryonic and central nervous system development^56^.

At the opposite extreme, six of the ten genic TRVs with the lowest *D* ratios occur in immune-related genes (Supplementary Table S4), including PRKCE, XRCC4, and CALCR, which have been implicated in innate immunity, antibody diversification, and immune-associated signaling, respectively^57–60^. One striking example exhibits nearly all unique alleles in the coding regions of PCGF3 (Fig. 3e), which acts primarily as a transcriptional activator required for mesodermal differentiation^61^, but also contributes to antiviral immunity by promoting interferon-responsive gene transcription^62^. Although extreme diversity compared to divergence alone could be caused by stabilizing selection coupled with high mutation rate^63^, the overrepresentation of immune genes is consistent with long-term balancing selection^64,65^.

We also consider genic TRVs with asymmetric *D* values, which reflect diversity in one species relative to the level of between-species divergence. Human-biased TRVs were enriched for cell morphogenesis and nervous system development terms, particularly neurogenesis (Extended Data Fig. 5a), and were primarily located in introns of cell signaling and intracellular trafficking genes, several with brain-specific activity (Supplementary Table S5). In contrast, chimp-biased TRVs were enriched only for cell-cell adhesion (Extended Data Fig. 5b), and occur largely in introns of genes involved in RNA processing, cell signaling and cytoskeletal organization (Supplementary Table S5). While such asymmetric *D* patterns can be explained by species-specific elevated mutation rates rapidly introducing new alleles, the genomic context of several of these TRVs suggests a contribution from directional selection, or a combination of both processes.

### Differential gene expression and TR variation

Given the predicted causal role of TRVs in gene expression variation in humans^1,8^, we analyzed 7,050 orthologous genes expressed in humans and chimpanzees across six tissues^66^. Genes harboring promoter TRVs did not exhibit greater expression divergence (Wilcoxon rank-sum test, FDR-adjusted p > 0.05 across all tissues; Supplementary Fig. 13). However, we identified a weak but significant positive correlation between absolute gene expression divergence and the average absolute TRV allele length divergence (Pearson’s r = 0.02, FDR-adjusted *p =* 0.006; Supplementary Fig. 14). This is consistent with studies suggesting that evolutionary changes in TR lengths can modulate expression levels of nearby genes^1,16^. Furthermore, this result agrees with studies showing that regions of elevated expression tend to exhibit an increase in the rate of mutations^67,68^, thus presumably harboring higher variation and divergence. When stratified by tissue type and genomic feature, significant positive correlations were observed in a subset of comparisons, including brain x intron TRs, kidney x 5’ UTR TRs, liver x intron TRs, and testis x promoter TRs (Supplementary Fig. 15). To further investigate this relationship, we focused on 4,125 differentially expressed (DE) genes between humans and chimpanzees in at least one of the six tissues, identified using *limma*^66^ (see Methods). While TRVs are generally not overrepresented in DE genes (Odds ratio = 0.84, *p =* 0.987), DE genes are significantly enriched for TRs with strong evidence to impact gene expression (eTRs^1,69^) (Odds ratio = 1.45, *p =* 0.012). Thus, while human-chimpanzee differential expression does not appear to be primarily driven by genomic TR species length variation, limited context-dependent associations may be due to a subset of regulatory TR.

Given longstanding evidence of the role of TRs in neuronal development and brain function^70,71^, we focused on a set of 738 DE genes between human and chimpanzee in organoids with telencephalon identity^20^. Again, we found a weak but significant positive correlation between mean absolute TR allele length divergence averaged over genes and absolute gene expression divergence (Supplementary Fig. 16a; Pearson’s r = 0.074, FDR-adjusted *p =* 0.0009). When stratified by cell type and genomic feature, positive correlations were significant for intermediate progenitor cells and radial glia cells in UTR and promoter TRVs (Fig. 4a; Extended Data Fig. 6). We also observe a modest but significant enrichment of TRs with larger mean lengths in humans among genes upregulated in humans relative to those upregulated in chimpanzees (Supplementary Fig. 16b; Odds ratio = 1.11, *p =* 1.05 × 10^−12^). These results suggest a connection between TR expansions and regulatory divergence in primates, in line with previous findings that TR divergence causes expression changes^19^.

**Fig. 4.**
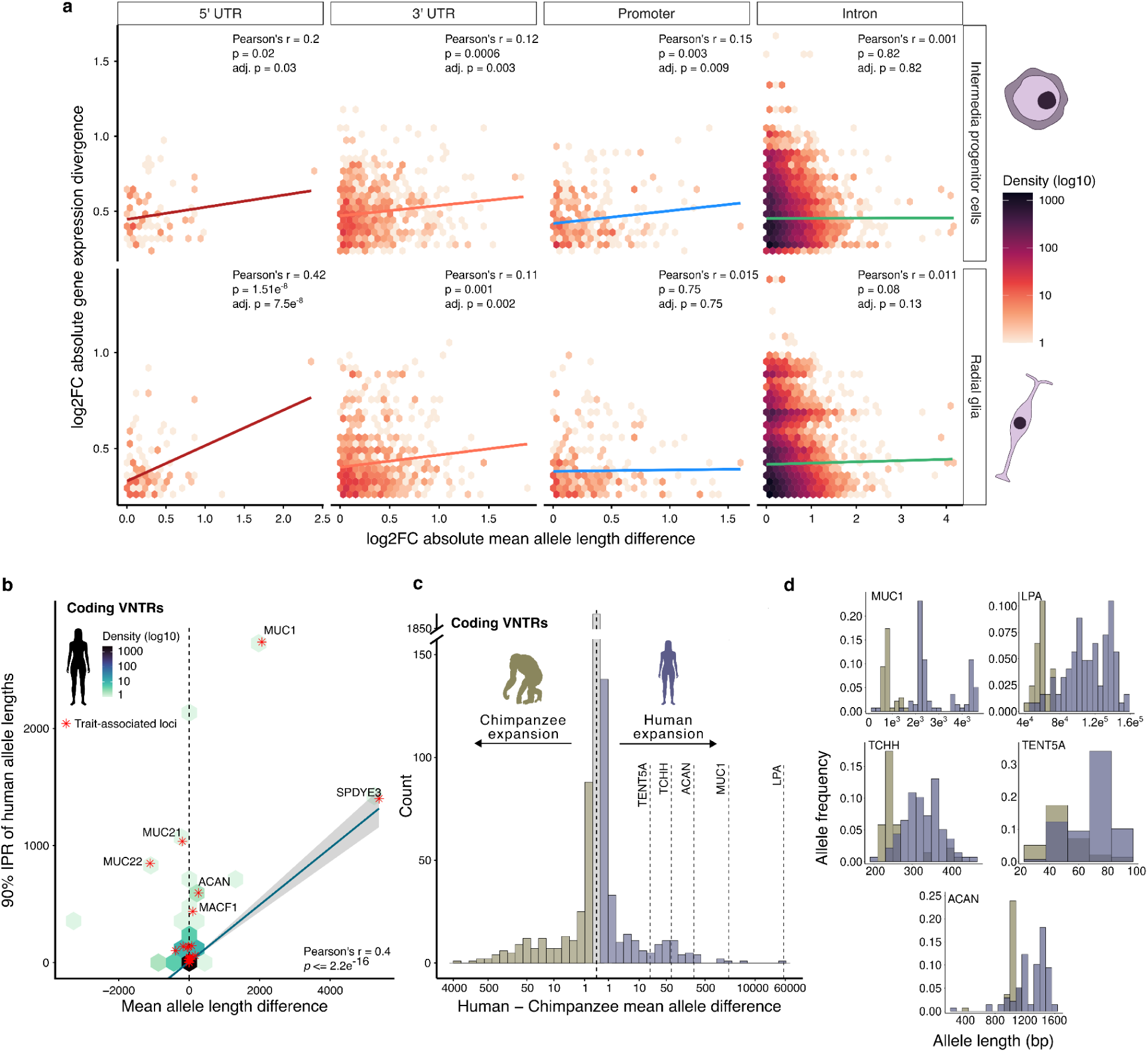
Gene expression, divergence, and trait-associated TRs. **a**, Heatmaps showing the correlation between absolute log fold change in mean TR allele length across TRs overlapping 5’UTR, 3’UTR and promoter, and intronic regions (x-axis) and absolute log fold change in gene expression divergence between humans and chimpanzee (y-axis) across intermediate progenitor cells and radial glia cells. **b**, TR mean allele length divergence between chimpanzees and humans (x-axis) versus the 90% interpercentile range of human allele lengths per locus. Red asterisks denote loci with evidence of genotype-phenotype association. The LPA locus is omitted. **c**, Distribution of mean allele length difference between humans and chimpanzees for coding VNTRs, highlighting five trait-causal VNTRs^74^. **d**, Chimpanzee and human allele frequency distributions for the same five loci.

By incorporating human and macaque cells from primary telencephalon samples into the DE analysis, Pollen et al., (2019) identified 261 candidate genes with human-specific regulatory changes during cortical development, which were significantly enriched for TRVs (Odds ratio = 2.15, *p =* 4.25 × 10^−6^). The majority of these candidate genes are up-regulated in humans (n=207) in one or more cell types, and several contain divergent TRVs (Supplementary Fig. 17). These patterns of gene expression divergence and allele length variation suggest that a subset of TRs might affect the regulation of genes involved in cortical neurogenesis, potentially contributing to human-specific aspects of neural development. Examples include VPS53 and PTPRS, which harbor highly divergent intronic VNTRs overlapping or near regulatory regions with high chromatin accessibility (Supplementary Fig. 18). Point mutations in VPS53 are linked to progressive cerebello-cerebral atrophy^72^ and increased levels of PTPRS caused by a point mutation are associated with decreased risk of Alzheimer’s disease^73^.

### Functional TR variation

Beyond their regulatory potential, TRs can directly affect protein structure and function. For example, a large-scale analysis of UK Biobank data identified a set of 25 coding VNTRs with significant trait association^74^. We found that of these 25 trait-associated TRs, 19 exhibit significant differences in length distribution between humans and chimpanzees, with 11 showing longer alleles in humans (Supplementary Fig. 19; Wilcoxon rank-sum test, FDR-adjusted p < 0.0001). Notably, nine of these fall within the top 2% of the ∼219,593 genic TRs showing higher mean lengths in humans, drawn from 837,612 homologous genic TRs.

These trait-associated VNTRs display not only high divergence but also increased human diversity (Fig. 4b). In a companion paper^23^, we show a similar trend in pathogenic TRs, which are consistently expanded and highly diverse in humans compared to chimpanzees. Notably, the 25 trait-associated VNTRs from Mukamel et al^74^ and 61 expansion-disorder TRs from the STRipy database^75^ are overrepresented among TRVs with exceptional relative length divergence and diversity in humans (top 10% in each)(Odds ratio = 10.66, CI: 5.13-20.51, *p =* 1.12 × 10^−8^). Focusing on five fine-mapped TR loci with strong evidence that length polymorphism, rather than linked SNVs or indels, directly drives trait association (posterior probability > 0.95^74^), we show that mean allele length difference between humans and chimpanzees was significantly greater than expected under the null distribution for all coding VNTRs (permutation test, 1,000,000 runs, *p* ≤ 1 × 10^−6^, and P = 0.003 when excluding the long and highly divergent VNTR in LPA) (Fig. 4c, d; Supplementary Fig. 20). Repeat length variation in these loci directly affects protein structure by altering the length of protein domains (see Fig. S1 from Mukamel et al.^74^). This pattern is expected for TRs with species-specific elevated mutability, particularly when longer expanded alleles have higher mutation rates^5,41^. Selection may also play a role if intermediate-length alleles are too weakly deleterious to overcome high mutation rates^76,77,78^, or if they are beneficial in an ecological context until further expansion causes fitness reduction, in the case of expansion disorders.

These observations support a hypothesis that highly expanded and diverse TRs are strong candidates for functional variation. This may be because increased TR length and variability provide a greater opportunity to alter genome structure and therefore function, potentially leading to a larger phenotypic impact^79,80^. In addition, by having disproportionately large trait effect sizes, the impact of these TRs may be more detectable, leading to ascertainment bias. Regardless of the underlying mechanism, our results suggest that these TRs are important targets for future large-scale comparative studies aimed at understanding the contribution of repeat variation to genome function, phenotypic diversity, and adaptation across evolutionary timescales.

## Discussion

TRs have long been hypothesized to play a significant role in phenotypic variation and evolutionary change. However, their evolutionary dynamics have remained underexplored due to limitations in resolving repeat variation at scale across species. Here, using assembly-level long-read data in a comparative framework, we present a comprehensive survey of the heterogeneous landscape of TR variation in primates, with focus on humans and chimpanzees. We recovered loci spanning a continuum of selective regimes, from pervasive length conservation in functional regions, consistent with stabilizing selection, to high divergence consistent with directional selection. We also observe loci with exceptional diversity, resulting either from balancing selection or elevated mutation rates and strong stabilizing selection, particularly outside functional or regulatory contexts.

TRs in CDS, and to a lesser extent 5’ UTRs, exhibit reduced polymorphism and extreme length conservation across evolutionary time (∼28.5 My; Fig. 1g, Fig. 2a-c), signatures of negative selection. However, while TRs are depleted in CDS, they are enriched in 5’ UTRs (Fig. 1f). This suggests that in these regions the high cost of their mutational risk is outweighed by a selected functional role, possibly in RNA folding^81^, translational regulation^82^, or modulation of transcription factor binding affinity^83^. Hence, TRs in 5’ UTR repeats are strong candidates for functional molecular impact.

We estimated TR-specific mutation rates based on TR divergence between humans and chimpanzees under a simple single-step mutation model (SMM) and selective neutrality, yielding results similar to those from trio-based average *de novo* mutation rate estimates^4^. This similarity suggests that, in broad strokes, on average the neutral SMM reasonably describes TR evolution, while groups of loci where the estimates depart suggest deviation from the SMM or neutrality. At the same time, we observe that species-specific expansions are associated with elevated within-species diversity, which could be caused by locus and species-specific increases in expansion-biased mutational processes.

Leveraging TR divergence-diversity ratios, we identified putative candidates for directional and balancing selection. TRs with both exceptional relative divergence and exceptional relative diversity were enriched in genes involved in nervous system development and synaptic processes, agreeing with prior associations between TR variation and neurodevelopment^8,71,84^. Notably, most known pathogenic repeat expansions are associated with neurological or neurodegenerative disorders, further supporting the particular sensitivity of nervous system processes to TR variation. We also identified TRVs with high diversity compared to divergence, including several extreme examples in immune-related genes. Although this pattern of variation may be due to high mutation rates coupled with stabilizing selection, it is also expected under long-term balancing selection, particularly at immune loci^21,85^.

We found associations between brain organoid expression divergence and divergence of TRs located in promoters and UTRs, where repeat variation can directly alter chromatin organization, nucleosome occupancy, and affect alternative splicing^16,86^. Further, genes with human-specific regulatory changes during cortical development are enriched in TRs, some of which are highly divergent. More broadly, trait and disease-causal TRs are associated bidirectionally with longer human alleles and higher diversity. These patterns may reflect elevated, lineage-specific mutational processes leading to runaway expansions with functional consequences, or possibly the action of directional selection shaping repeat length distributions. While the underlying mechanism remains unresolved, we hypothesize that exceptional length divergence and species-specific diversity may serve as useful criteria for prioritizing TRs for functional studies.

While we provide a comprehensive overview of TR variation in primates, several limitations should be noted. First, estimates of within-species diversity are restricted to humans and chimpanzees. Second, we focused on loci shared between species. Future studies including species-specific TRs may provide additional insight into forces impacting TR birth and death. Third, current TR-phenotype association evidence is largely derived from human short-reads, which have limited resolution for long TRs and do not capture trait associations in other species. Comparable long-read studies across NHP are essential to assess whether similar patterns in trait-causal loci extend beyond humans. Finally, our analyses are limited to TR allele length polymorphism and do not capture other forms of variation capable of affecting phenotypes, such as sequence interruptions and changes in motif composition^87^.

In summary, our results show that TR variation reflects the interplay of mutational processes and selective pressures, acting both as conserved functional elements and as fuel for evolutionary innovation. Ongoing advances in TR genotyping and mutation inference, along with increasingly accessible population-scale cross-species variant catalogs, will enable systematic characterization of TR mutational dynamics and their contribution to adaptation across evolutionary timescales.

## Methods

### TR reference catalogs

Using the TRACK pipeline^88^, we generated TR catalogs for eight ape T2T reference genomes^6,89^ (Fig. 1b). Briefly, TRs were identified with Tandem Repeat Finder v.4.09^90^ using the following parameters: *matchscore* 2, *mismatchscore* 5, *indelscore* 7, *pm* 80, *pi* 10, *minscore* 24, *maxperiod* 2000, -*l* 6. Resulting annotations were filtered by total length (>11 bp and <10 Kbp), copy number (>2.5), and constancy score, i.e., sequence similarity between adjacent repeat units (>60%). When TRs overlapped by 5 bp or less, we retained the repeat with the shortest motif length. We then normalized motif sequences to their smallest periodic units. For example, the motif “ATATAT” was reduced to “AT”, and the copy number was recalculated accordingly. This normalization step was implemented because TRF reports consensus motifs that maximize alignment scores, which do not always correspond to the minimal motifs. As a result, TRF output may contain nested motifs composed of repeated instances of a shorter underlying unit. Finally, we queried our catalog against the Dfam database to identify instances where TRs intersect known repetitive element families^91^. After this initial characterization of the catalogs, we applied additional filtering to exclude centromeric regions and regions containing alpha satellite DNA (cenSat) and subterminal satellites (StSat). Annotations were obtained from the CHM13 and T2T-Primate Consortium to identify and remove complex, high-order repeat (HOR)-rich regions before homology assessment.

### Annotation of TR genomic features

To classify TRs by genomic feature, we used the GENCODE GFF2 gene annotation for CHM13. We retained only transcripts annotated as the APPRIS principal isoform to ensure high-confidence gene models^92^. Exons, coding sequences (CDS), and introns were directly extracted from the annotation. Transcription start sites (TSS) were defined as the 5’ end of each transcript, and promoter regions were defined as the 1 Kb upstream of the TSS, accounting for strand orientation. Untranslated regions (UTRs) were inferred based on exon coordinates: 5’ UTRs were inferred as the exon segments upstream of the start codon, and 3’ UTRs as those downstream of the stop codon, considering strand orientation. To avoid overestimating UTRs, we required inferred regions to not overlap CDS regions. The TR catalog was intersected to these annotations and a hierarchical classification was applied to resolve edge cases where TRs overlap more than one feature: CDS > 5’ UTR > 3’ UTR > promoter > intron > intergenic.

### Homology assessment

To identify homologous TRs between reference genomes, we used a multi-step alignment-based pipeline implemented in TRACK^88^. TR coordinates from a target genome were lifted to a query genome using the UCSC LiftOver tool^93^ with *-minMatch* 0.1 and *-bedPlus=3 -tab* to preserve metadata tracking of TRs across genome builds. Lifted TRs were intersected with the native TR catalog of the query genome using *bedtools intersect*, requiring a reciprocal overlap of at least 10%.

For each overlapping TR pair, we extracted and compared their motifs. To ensure strand-invariant and phase-independent comparison, TRACK computes the lexicographically smallest cyclic permutation of each motif and its reverse complement. This step allows the direct comparison of motif sequences regardless of strand orientation or rotational phase. To assess motif similarity, we performed global pairwise alignments between each candidate homologous TR pair with EMBOSS Needle^94^, using the following parameters: -*gapopen 10* and *-gapextend 0.5*. Each motif pair was aligned in both forward and reverse-complement orientation, and we retained the alignment with the highest similarity score. TR pairs with the best alignment similarity score >=95% were retained as confidently homologous. Homology detection was performed bidirectionally, with both genomes in the pair used as target and query. The final set of homologous TRs was defined as the intersection of high-similarity TR pairs identified in both directions. This reciprocal filtering strategy ensures that homologs are robust to mapping artifacts or asymmetric genome annotations.

To quantify the correlation of normalized TR density between species we computed density in non-overlapping 1 Mb windows in the human genome, and windows were lifted over to the corresponding NHP genomes. Lifted windows were filtered based on length, retaining regions between 0.8-2 Mb for most species, and 0.2-2 Mb for the more divergent taxa, i.e. siamang gibbon and macaque, where liftover performance is reduced. For each species, TR density was calculated as the proportion of bps covered by TRs within each lifted genomic window and normalized to sum to 1 prior to comparison.

### Determining TR genotypes

TRs were genotyped using PacBio HiFi data from 46 humans in the Human Pangenome Research Consortium (HPRC)^95^ and 23 near-T2T chimpanzee genomes, described in a companion paper^23^, with Tandem Repeat Genotyping Tool v.3.0 (TRGT)^96^. Variants were filtered for missing data (*--max-missing* 1), minimum allele spanning depth (>3) and allele constancy score (>0.6).

### Estimating TR mutation rates

Per-locus mutation rates were estimated under the stepwise mutation model (SMM) and neutrality based on the observed genetic distance between humans and chimpanzees. Specifically, we computed the TR genetic distance, originally termed δµ^2^, but we refer to as *d*^45^:

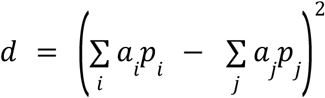

where *a_i_* is allele copy number and *p_i_* is the allele frequency for allele *i* in humans, and *j* indexed quantities in chimpanzees. Under neutrality and the SMM, the expected value of *d* increases linearly with the mutation rate (μ) and the divergence time (*t*)^45^. We therefore estimated per-generation mutation rates as

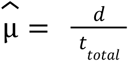

where *t_total_* is the sum of the human and chimpanzee branch lengths. We assumed branch lengths of 7.7 million years for each lineage and generation times of 29 years for humans and 25 years for chimpanzees, yielding *t*_*total*_ = 5. 8 × 10^5^generations^97^.

### Divergence-diversity ratio (*D*)

TRV per locus divergence-diversity ratio (*D*) was computed as the ratio of between-species to within-species variance in allele length, computed separately for each species after removing 3 outlier alleles per species:

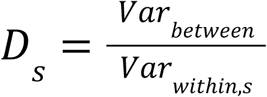

where between-species variance for each TR was defined as the weighted squared difference between each species-specific mean length and the overall mean length across both species:

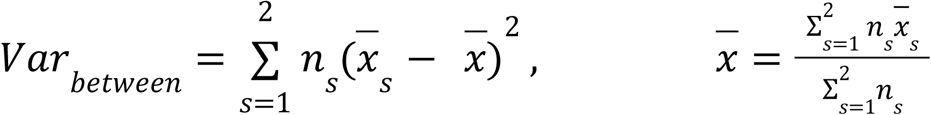

where *n*_s_ is the number of alleles in species *s,* and *x* is the overall mean length across both species. One characteristic of *D*_s_ is that elevated within-species diversity can also inflate between species variance, reducing its power to detect loci with exceptionally high relative diversity.

TRs with high *D_s_* in both species were defined as the top 1,000 loci in both species, while TRs with low *D_s_* in both species were defined as the bottom 1,000 loci after excluding repeats with within-species variance below 1. We implemented this threshold as a conservative noise floor in order to eliminate small TRs with low variance that produce small ratios likely not biologically relevant (Supplementary Fig. 1). Species-specific extremes were defined based on deviation from the diagonal in the human-chimpanzee *D* comparison, representing loci with pronounced divergence relative to diversity in a single lineage.

### Gene ontology enrichment analysis

We first performed Gene Ontology (GO) enrichment analysis to test whether genes containing TRs were associated with specific biological process terms, using the full set of human genes as a background set. Next, we selected genes intersecting the top 1,000 TRVs within each group of extreme *D* values, using the full set of genes intersecting TRs as the background set. Analyses were conducted in ShinyGO v.0.82^98^ with a minimum pathway size of 15 and a maximum pathway size of 1,000 genes. Significance was assessed using the false discovery rate (FDR), and only pathways with FDR-corrected *p* < 0.01 were considered enriched. To increase confidence in the results, we performed enrichment analysis with 10 random sets of 1,000 TR-containing genes, which showed no GO enrichment.

### TR divergence and gene expression divergence

Gene expression data were obtained from Brawand et al.^66^, which provides normalized RPKM (reads per kilobase of exon per million mapped reads) values for six tissues (brain, cerebellum, heart, kidney, liver, and testis) from humans and chimpanzees. To reduce noise from lowly expressed genes, we applied an RPKM threshold of >1 in at least one individual for each tissue and species. Differential expression (DE) between species was assessed per tissue using *limma*^99^ on log-transformed RPKM values, with empirical Bayes moderation of gene-wise standard errors. Genes with FDR-adjusted p-values < 0.05 were considered DE. TR length divergence was quantified as the log2 fold change of mean allele length per locus and compared to gene expression divergence using Pearson’s correlation. To focus on genes containing TRs known to be associated with expression (eTRs), we retained those associated with fine-mapped expression STR^1^ and expression-associated VNTRs^69^.

## Code Availability

All code used in the paper can be obtained at https://github.com/caroladam/tr_analyses.

## Data availability

The HPRC data can be obtained at https://humanpangenome.org/data/. The chimpanzee assemblies can be obtained at [pending]. Chimpanzee-human homologous TR catalog and VCF files for humans and chimpanzees can be obtained at 10.5281/zenodo.20616249 [pending publication].

## Acknowledgements

We thank the Rohlfs lab for helpful discussion.

## Funding statement

This work was supported by the National Science Foundation (NSF) CAREER award 2144878 to R.V.R., the NIH National Institute of General Medicine award R35GM142916 to P.H.S., and the NIH National Human Genome Research Institute award R01HG013017 to P.H.S.

## Author contributions

R.V.R., P.H.S. conceived the design of the study, acquired funding, and supervised the research. C.L.A, J.L.R., R.V.R., P.H.S. processed and analyzed the data. C.L.A, R.R. drafted the manuscript. C.L.A, J.L.R., R.V.R., P.H.S. reviewed and edited the manuscript.

## Competing interests

The authors declare no competing financial interests.

## Additional information

Supplementary information is available for this paper. Correspondence should be addressed to R.V.R. and C.L.A.

---

**Extended Data Fig. 1.**
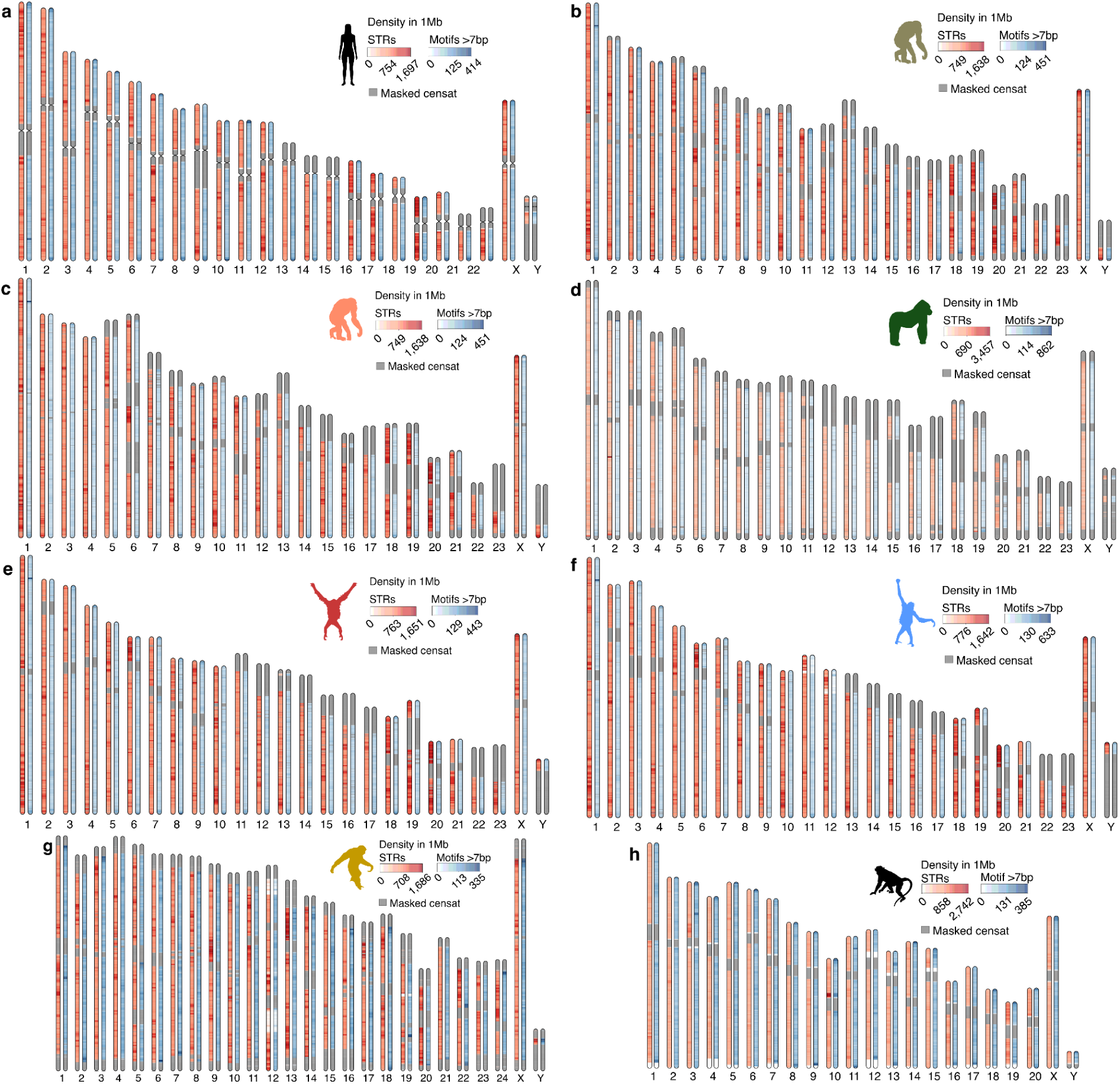
Genomic TR distributions in non-human primate genomes. Ideograms showing the density of STRs in red and VNTRs in blue across non-overlapping 1 Mb windows. Masked CenSat regions are marked in gray. TR densities are shown for **a**, *Homo sapiens*; **b**, *Pan troglodytes*; **c**, *Pan paniscus*; **d**, *Gorilla gorilla*; **e**, *Pongo pygmaeus*; **f**, *Pongo abelii*; **g**, *Symphalangus syndactylus*; and **h**, *Macaca mulatta*.

**Extended Data Fig. 2.**
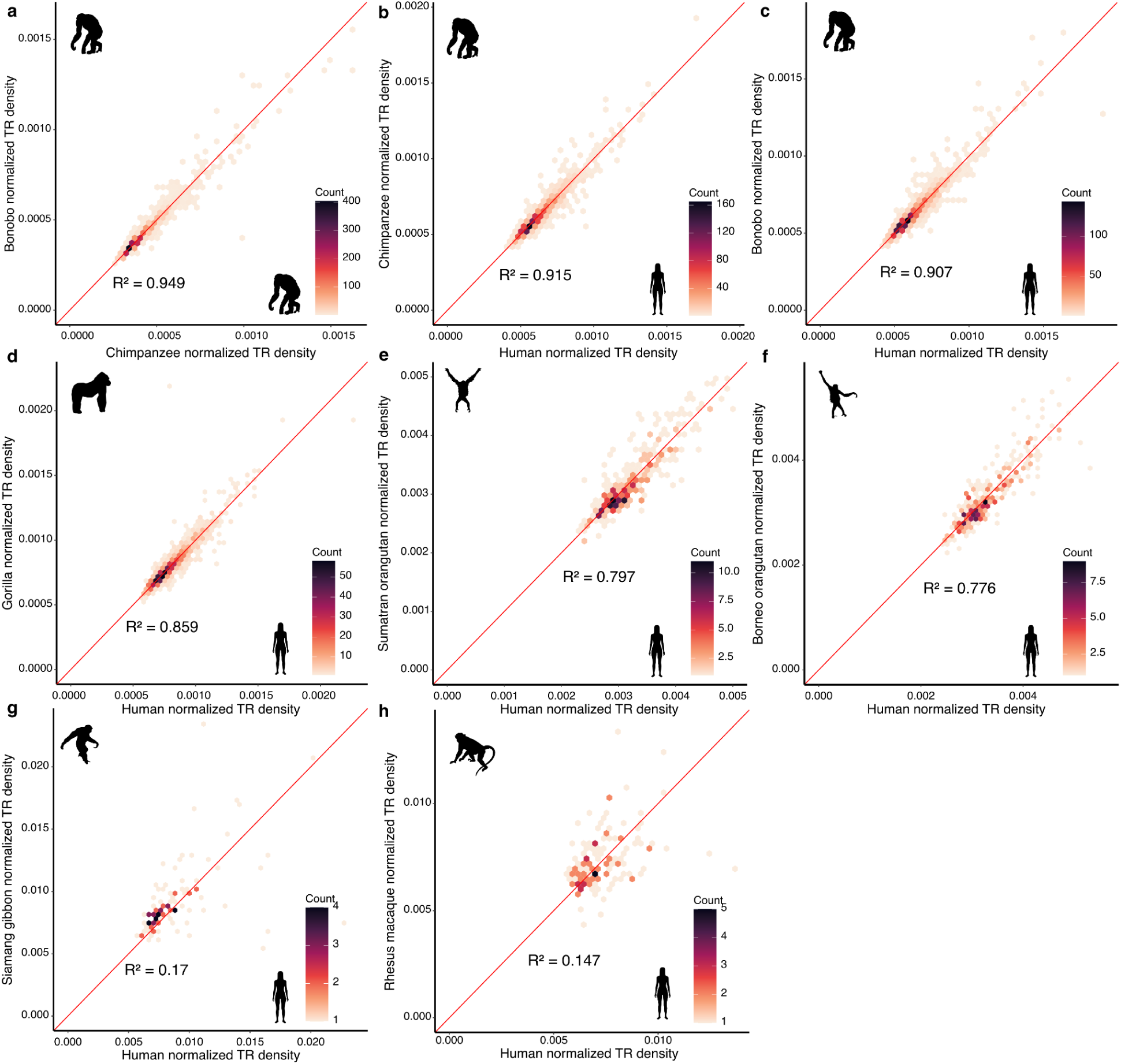
Correlation of TR density in homologous regions. Correlation between normalized density of TRs between **a**, chimpanzee and bonobo; and between human and **b**, chimpanzee; **c**, bonobo; **d**, gorilla; **e**, Sumatran orangutan; **f**, Borneo orangutan; **g**, siamang gibbon; and **h**, macaque.

**Extended Data Fig. 3.**
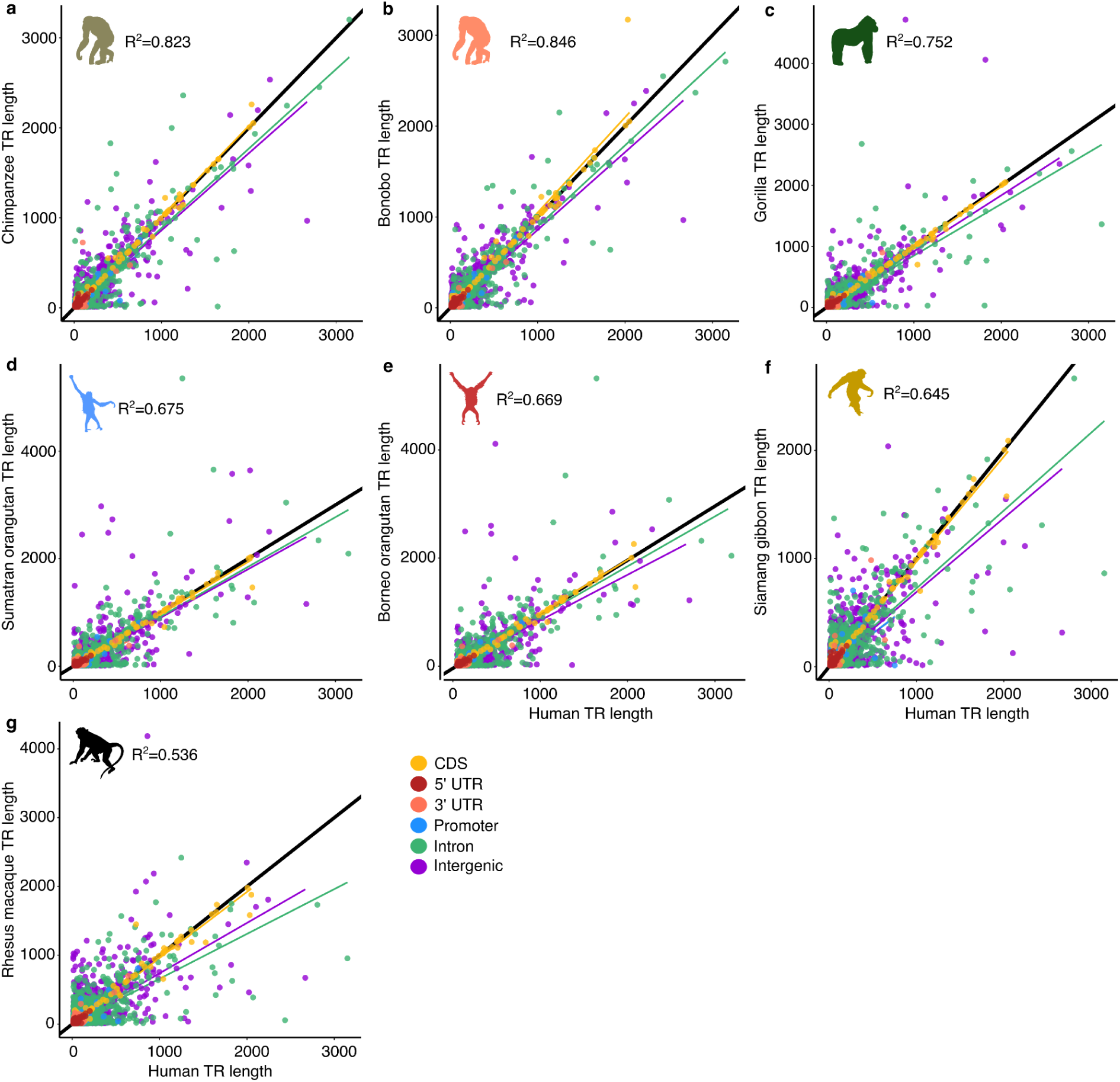
Landscape of TR length variation across human and non-human primate T2T reference genomes. Scatterplot of reference TR lengths between human (x-axis) and seven non-human T2T reference primates (y-axis): **a**, *Pan troglodytes*; **b**, *Pan paniscus*; **c**, *Gorilla gorilla*; **d**, *Pongo pygmaeus*; **e**, *Pongo abelii*; **f**, *Symphalangus syndactylus;* and **g**, *Macaca mulatta*. Each point represents a single TR locus, color-coded according to CHM13 genomic annotation.

**Extended Data Fig. 4.**
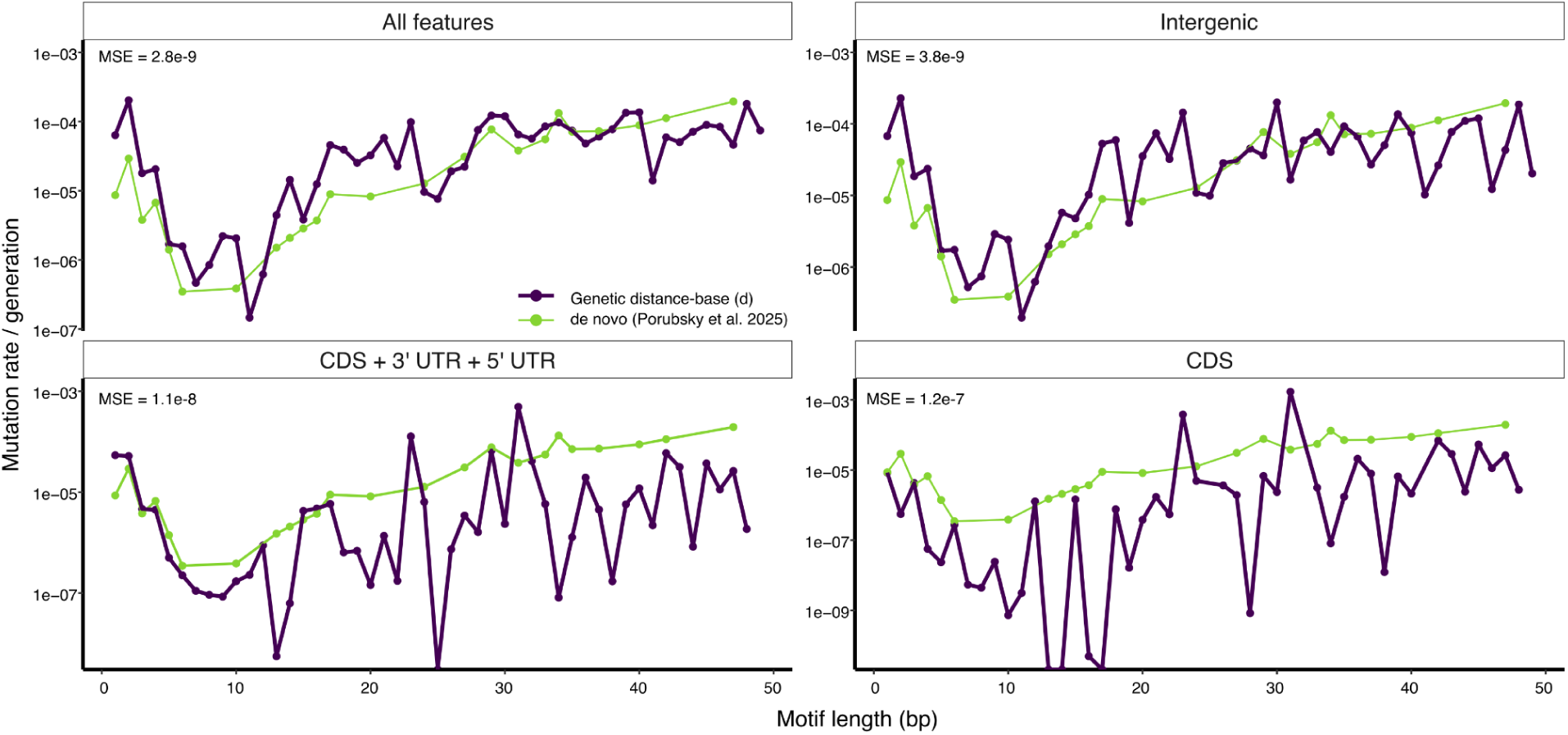
Estimates of *de novo* and genetic distance-based (*d*) TR mutation rates averaged across motif lengths for different subsets of loci.

**Extended Data Fig. 5.**
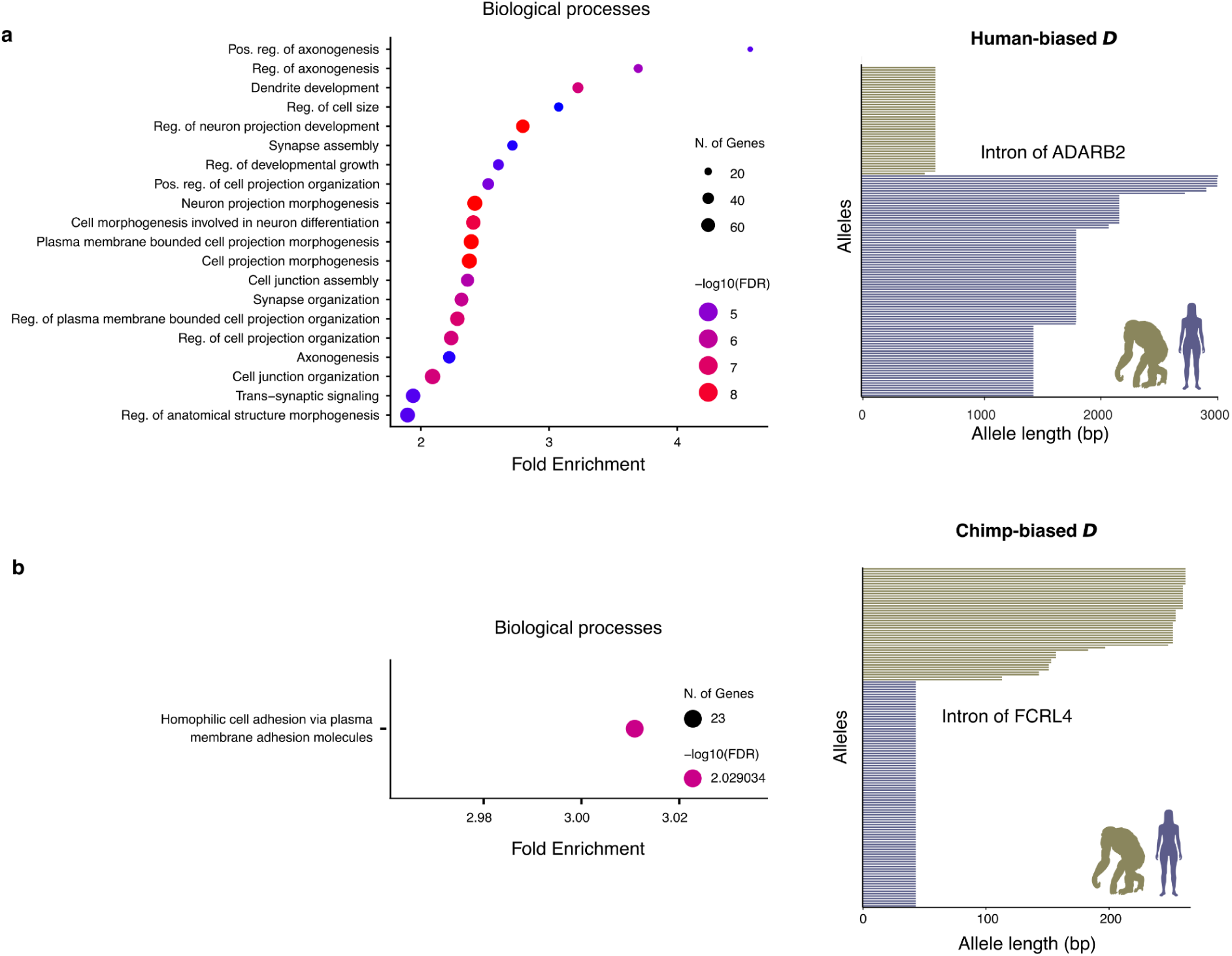
Biological processes associated with asymmetric divergence-diversity (*D*) ratios. Gene Ontology enrichment analysis for biological processes terms associated with genes intersecting the top 1,000 genic TRs with asymmetric *D*, and allele length distribution of one representative genic TR with **a**, human-biased *D*, with an intronic TR at ADARB2; and **b**, chimp-biased *D*, with an intronic TR at FCRL4.

**Extended Data Fig. 6.**
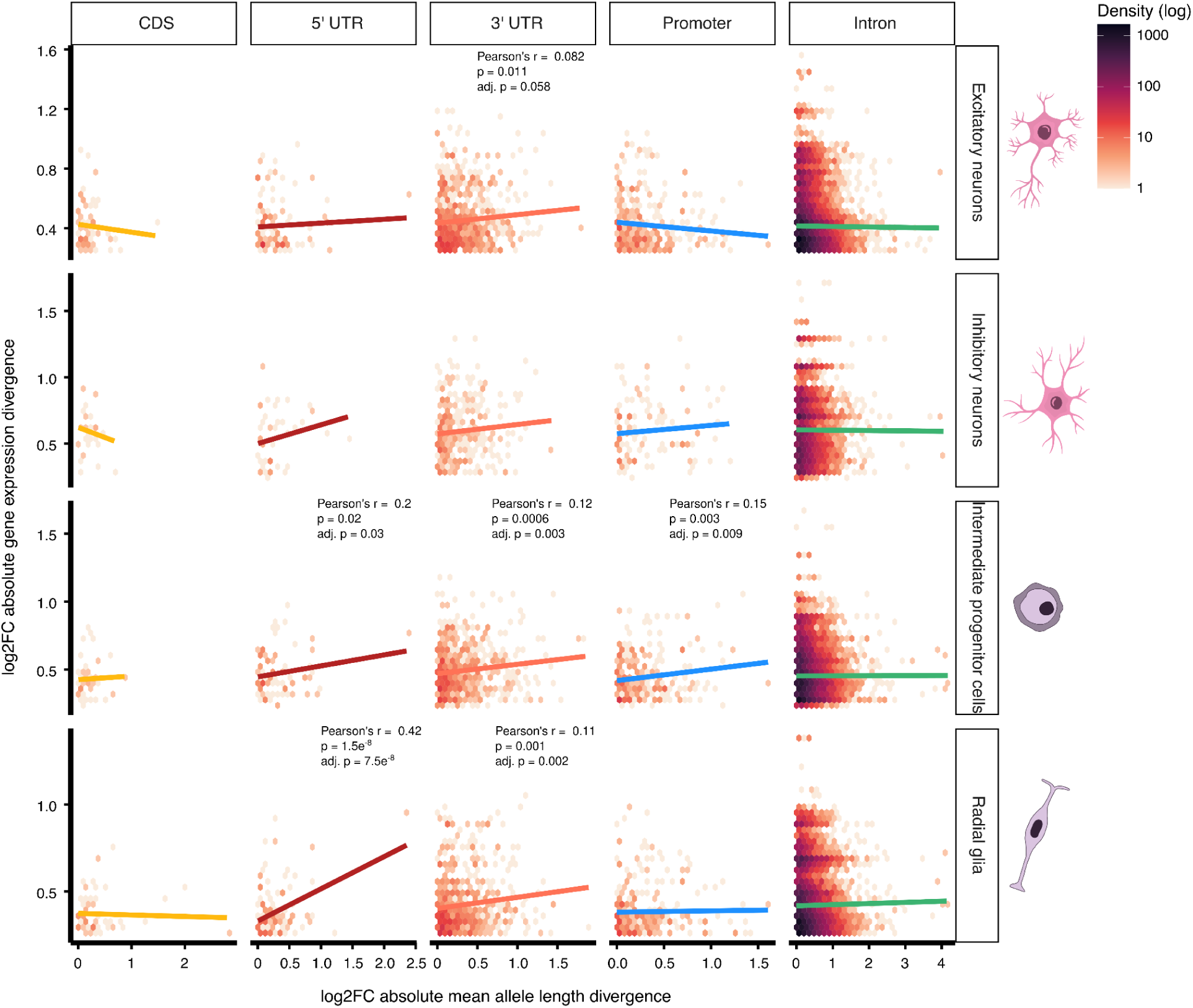
Heatmaps showing the correlation between absolute log fold change in mean TR allele length across genes in each genomic feature (x-axis) and absolute log fold change in gene expression divergence between humans and chimpanzees (y-axis) across organoids with telencephalon identity from Pollen et al. (2019).

## Notes

https://doi.org/10.5281/zenodo.20616248

### Competing Interest Statement

The authors have declared no competing interest.

### Summary of Updates

This version provides added analyses, updated figures and updated supplementary material. Major modifications included analyses of the association between TR divergence and differential expression between humans and chimpanzees, and analyses of functional TRs.

## References

1. Fotsing, S. F. et al. The impact of short tandem repeat variation on gene expression. Nat. Genet. 51, 1652–1659 (2019).

2. Doyle, L. et al. Rational design of α-helical tandem repeat proteins with closed architectures. Nature 528, 585–588 (2015).

3. Hannan, A. J. Tandem repeat polymorphisms: modulators of disease susceptibility and candidates for ‘missing heritability’. Trends Genet. 26, 59–65 (2010).

4. Porubsky, D. et al. Human de novo mutation rates from a four-generation pedigree reference. Nature 643, 427–436 (2025).

5. Brinkmann, B., Klintschar, M., Neuhuber, F., Hühne, J. & Rolf, B. Mutation rate in human microsatellites: influence of the structure and length of the tandem repeat. Am. J. Hum. Genet. 62, 1408–1415 (1998).

6. Nurk, S. et al. The complete sequence of a human genome. Science 376, 44–53 (2022).

7. Fan, H. & Chu, J.-Y. A brief review of short tandem repeat mutation. Genomics Proteomics Bioinformatics 5, 7–14 (2007).

8. Gymrek, M. et al. Abundant contribution of short tandem repeats to gene expression variation in humans. Nat. Genet. 48, 22–29 (2016).

9. Erwin, G. S. et al. Recurrent repeat expansions in human cancer genomes. Nature 613, 96–102 (2023).

10. Schloissnig, S., et al. Long-read sequencing and structural variant characterization in 1,019 samples from the 1000 Genomes Project. bioRxivorg (2024) doi:10.1101/2024.04.18.590093.

11. Song, J. H. T., Lowe, C. B. & Kingsley, D. M. Characterization of a human-specific tandem repeat associated with bipolar disorder and schizophrenia. Am. J. Hum. Genet. 103, 421–430 (2018).

12. Huang, Y. et al. Short tandem repeats in populations of the Qinghai-Tibet Plateau and adjacent regions provide insights into high-altitude adaptation. Sci. Adv. 11, eadx1590 (2025).

13. Zhou, K., Aertsen, A. & Michiels, C. W. The role of variable DNA tandem repeats in bacterial adaptation. FEMS Microbiol. Rev. 38, 119–141 (2014).

14. Fondon, J. W., 3rd & Garner, H. R. Molecular origins of rapid and continuous morphological evolution. Proc. Natl. Acad. Sci. U. S. A. 101, 18058–18063 (2004).

15. Villanea, F. A. et al. The MUC19 gene: An evolutionary history of recurrent introgression and natural selection. Science 389, eadl0882 (2025).

16. Vinces, M. D., Legendre, M., Caldara, M., Hagihara, M. & Verstrepen, K. J. Unstable tandem repeats in promoters confer transcriptional evolvability. Science 324, 1213–1216 (2009).

17. L Rocha, J., Lou, R. N. & Sudmant, P. H. Structural variation in humans and our primate kin in the era of telomere-to-telomere genomes and pangenomics. Curr. Opin. Genet. Dev. 87, 102233 (2024).

18. Liu, Q. & Tian, W. Association of human-specific expanded short tandem repeats with neuron-specific regulatory features. Sci. Adv. 11, eadp9707 (2025).

19. Sulovari, A. et al. Human-specific tandem repeat expansion and differential gene expression during primate evolution. Proc. Natl. Acad. Sci. U. S. A. 116, 23243–23253 (2019).

20. Pollen, A. A. et al. Establishing cerebral organoids as models of human-specific brain evolution. Cell 176, 743–756.e17 (2019).

21. Leffler, E. M. et al. Multiple instances of ancient balancing selection shared between humans and chimpanzees. Science 339, 1578–1582 (2013).

22. Prado-Martinez, J. et al. Great ape genetic diversity and population history. Nature 499, 471–475 (2013).

23. Rocha, J., et al. A Pan-pangenome illuminates complex structural variation and selection in humans, chimpanzees, and bonobos. bioRxiv (2026) doi:10.64898/2026.06.06.730619.

24. Sharma, A. & Sowpati, D. T. Analysis of tandem repeats in seven telomere-to-telomere primate genomes. J. Genet. 104, 14 (2025).

25. Srivastava, S., Avvaru, A. K., Sowpati, D. T. & Mishra, R. K. Patterns of microsatellite distribution across eukaryotic genomes. BMC Genomics 20, 153 (2019).

26. Verbiest, M. et al. Mutation and selection processes regulating short tandem repeats give rise to genetic and phenotypic diversity across species. J. Evol. Biol. 36, 321–336 (2023).

27. El-Sawy, M. & Deininger, P. Tandem insertions of Alu elements. Cytogenet. Genome Res. 108, 58–62 (2005).

28. Webster, M. T., Smith, N. G. C. & Ellegren, H. Microsatellite evolution inferred from human-chimpanzee genomic sequence alignments. Proc. Natl. Acad. Sci. U. S. A. 99, 8748–8753 (2002).

29. Grimwood, J. et al. The DNA sequence and biology of human chromosome 19. Nature 428, 529–535 (2004).

30. Subramanian, S., Mishra, R. K. & Singh, L. Genome-wide analysis of microsatellite repeats in humans: their abundance and density in specific genomic regions. Genome Biol. 4, R13 (2003).

31. Rhie, A. et al. The complete sequence of a human Y chromosome. Nature 621, 344–354 (2023).

32. Linthorst, J. et al. Extreme enrichment of VNTR-associated polymorphicity in human subtelomeres: genes with most VNTRs are predominantly expressed in the brain. Transl. Psychiatry 10, 369 (2020).

33. Cechova, M. et al. High satellite repeat turnover in great apes studied with short- and long-read technologies. Mol. Biol. Evol. 36, 2415–2431 (2019).

34. Koga, A., Hirai, Y., Hara, T. & Hirai, H. Repetitive sequences originating from the centromere constitute large-scale heterochromatin in the telomere region in the siamang, a small ape. Heredity (Edinb.) 109, 180–187 (2012).

35. Madsen, B. E., Villesen, P. & Wiuf, C. Short tandem repeats in human exons: a target for disease mutations. BMC Genomics 9, 410 (2008).

36. Usdin, K., House, N. C. M. & Freudenreich, C. H. Repeat instability during DNA repair: Insights from model systems. Crit. Rev. Biochem. Mol. Biol. 50, 142–167 (2015).

37. Schaper, E., Gascuel, O. & Anisimova, M. Deep conservation of human protein tandem repeats within the eukaryotes. Mol. Biol. Evol. 31, 1132–1148 (2014).

38. Byeon, G. W. et al. Functional and structural basis of extreme conservation in vertebrate 5’ untranslated regions. Nat. Genet. 53, 729–741 (2021).

39. Wieder, N. et al. The role of untranslated region variants in Mendelian disease: a review. Eur. J. Hum. Genet. 33, 1096–1105 (2025).

40. Kronenberg, Z. N. et al. High-resolution comparative analysis of great ape genomes. Science 360, eaar6343 (2018).

41. Neff, B. D. & Gross, M. R. Microsatellite evolution in vertebrates: inference from AC dinucleotide repeats. Evolution 55, 1717–1733 (2001).

42. Bilgin Sonay, T., et al. Tandem repeat variation in human and great ape populations and its impact on gene expression divergence. Genome Res. 25, 1591–1599 (2015).

43. Li, W. H. & Sadler, L. A. Low nucleotide diversity in man. Genetics 129, 513–523 (1991).

44. Willems, T. et al. The landscape of human STR variation. Genome Res. 24, 1894–1904 (2014).

45. Goldstein, D. B., Ruiz Linares, A., Cavalli-Sforza, L. L. & Feldman, M. W. Genetic absolute dating based on microsatellites and the origin of modern humans. Proc. Natl. Acad. Sci. U. S. A. 92, 6723–6727 (1995).

46. Gymrek, M., Willems, T., Reich, D. & Erlich, Y. Interpreting short tandem repeat variations in humans using mutational constraint. Nat. Genet. 49, 1495–1501 (2017).

47. Sainudiin, R., Durrett, R. T., Aquadro, C. F. & Nielsen, R. Microsatellite mutation models: insights from a comparison of humans and chimpanzees. Genetics 168, 383–395 (2004).

48. Mitra, I. et al. Patterns of de novo tandem repeat mutations and their role in autism. Nature 589, 246–250 (2021).

49. Lai, Y. & Sun, F. The relationship between microsatellite slippage mutation rate and the number of repeat units. Mol. Biol. Evol. 20, 2123–2131 (2003).

50. Beaumont, M. A. & Balding, D. J. Identifying adaptive genetic divergence among populations from genome scans. Mol. Ecol. 13, 969–980 (2004).

51. Excoffier, L., Foll, M. & Petit, R. J. Genetic consequences of range expansions. Annu. Rev. Ecol. Evol. Syst. 40, 481–501 (2009).

52. Ellegren, H. Microsatellites: simple sequences with complex evolution. Nat. Rev. Genet. 5, 435–445 (2004).

53. Cassandri, M. et al. Zinc-finger proteins in health and disease. Cell Death Discov. 3, 17071 (2017).

54. Jovanovic, V. M. et al. Positive selection in gene regulatory factors suggests adaptive pleiotropic changes during human evolution. Front. Genet. 12, 662239 (2021).

55. Nowick, K. et al. Gain, loss and divergence in primate zinc-finger genes: a rich resource for evolution of gene regulatory differences between species. PLoS One 6, e21553 (2011).

56. Elsir, T., Smits, A., Lindström, M. S. & Nistér, M. Transcription factor PROX1: its role in development and cancer. Cancer Metastasis Rev. 31, 793–805 (2012).

57. Altman, A. & Kong, K.-F. Protein kinase C enzymes in the hematopoietic and immune systems. Annu. Rev. Immunol. 34, 511–538 (2016).

58. Soulas-Sprauel, P. et al. Role for DNA repair factor XRCC4 in immunoglobulin class switch recombination. J. Exp. Med. 204, 1717–1727 (2007).

59. Wang, S., Wang, W. & Zeng, J. Role of CALCR expression in liver cancer: Implications for the immunotherapy response. Mol. Med. Rep. 31, 41 (2025).

60. Maleitzke, T. et al. The calcitonin receptor protects against bone loss and excessive inflammation in collagen antibody-induced arthritis. iScience 25, 103689 (2022).

61. Zhao, W. et al. Polycomb group RING finger proteins 3/5 activate transcription via an interaction with the pluripotency factor Tex10 in embryonic stem cells. J. Biol. Chem. 292, 21527–21537 (2017).

62. Da, G. et al. Nuclear PCGF3 inhibits the antiviral immune response by suppressing the interferon-stimulated gene. Cell Death Discov. 10, 429 (2024).

63. Zhang, X.-S. & Hill, W. G. Genetic variability under mutation selection balance. Trends Ecol. Evol. 20, 468–470 (2005).

64. Bitarello, B. D. et al. Signatures of long-term balancing selection in human genomes. Genome Biol. Evol. 10, 939–955 (2018).

65. Minias, P. & Vinkler, M. Selection balancing at innate immune genes: Adaptive polymorphism maintenance in Toll-like receptors. Mol. Biol. Evol. 39, msac102 (2022).

66. Brawand, D. et al. The evolution of gene expression levels in mammalian organs. Nature 478, 343–348 (2011).

67. Park, C., Qian, W. & Zhang, J. Genomic evidence for elevated mutation rates in highly expressed genes. EMBO Rep. 13, 1123–1129 (2012).

68. Bachl, J., Carlson, C., Gray-Schopfer, V., Dessing, M. & Olsson, C. Increased transcription levels induce higher mutation rates in a hypermutating cell line. J. Immunol. 166, 5051–5057 (2001).

69. Eslami Rasekh, M., Hernández, Y., Drinan, S. D., Fuxman Bass, J. I. & Benson, G. Genome-wide characterization of human minisatellite VNTRs: population-specific alleles and gene expression differences. Nucleic Acids Res. 49, 4308–4324 (2021).

70. Hammock, E. A. D. & Young, L. J. Microsatellite instability generates diversity in brain and sociobehavioral traits. Science 308, 1630–1634 (2005).

71. Xiao, X. et al. Revisiting tandem repeats in psychiatric disorders from perspectives of genetics, physiology, and brain evolution. Mol. Psychiatry 27, 466–475 (2022).

72. Feinstein, M. et al. VPS53 mutations cause progressive cerebello-cerebral atrophy type 2 (PCCA2). J. Med. Genet. 51, 303–308 (2014).

73. Poirier, A. et al. PTPRS is a novel marker for early Tau pathology and synaptic integrity in Alzheimer’s disease. Sci. Rep. 14, 14718 (2024).

74. Mukamel, R. E. et al. Protein-coding repeat polymorphisms strongly shape diverse human phenotypes. Science 373, 1499–1505 (2021).

75. Halman, A., Dolzhenko, E. & Oshlack, A. STRipy: A graphical application for enhanced genotyping of pathogenic short tandem repeats in sequencing data. Hum. Mutat. 43, 859–868 (2022).

76. Pajic, P. & Gokcumen, O. Evolutionary balancing of genetic consequence and innovation in mammals through variable number tandem repeats. Genome Biol. Evol. 18, evaf250 (2026).

77. Press, M. O., Hall, A. N., Morton, E. A. & Queitsch, C. Substitutions are boring: Some arguments about parallel mutations and high mutation rates. Trends Genet. 35, 253–264 (2019).

78. Ibañez, K. et al. Increased frequency of repeat expansion mutations across different populations. Nat. Med. 30, 3357–3368 (2024).

79. Gemayel, R., Vinces, M. D., Legendre, M. & Verstrepen, K. J. Variable tandem repeats accelerate evolution of coding and regulatory sequences. Annu. Rev. Genet. 44, 445–477 (2010).

80. Manigbas, C. A. et al. A phenome-wide association study of tandem repeat variation in 168,554 individuals from the UK Biobank. Nat. Commun. 15, 10521 (2024).

81. Kinney, N., Pathak, D., Evans, E. & Arias, P. Short tandem repeat variants are possibly associated with RNA secondary structure and gene expression. PLoS One 20, e0326355 (2025).

82. Leppek, K., Das, R. & Barna, M. Functional 5’ UTR mRNA structures in eukaryotic translation regulation and how to find them. Nat. Rev. Mol. Cell Biol. 19, 158–174 (2018).

83. Horton, C. A. et al. Short tandem repeats bind transcription factors to tune eukaryotic gene expression. Science 381, eadd1250 (2023).

84. Cui, Y. et al. Multi-omic quantitative trait loci link tandem repeat size variation to gene regulation in human brain. Nat. Genet. 57, 369–378 (2025).

85. Ferrer-Admetlla, A. et al. Balancing selection is the main force shaping the evolution of innate immunity genes. J. Immunol. 181, 1315–1322 (2008).

86. Hamanaka, K. et al. Genome-wide identification of tandem repeats associated with splicing variation across 49 tissues in humans. Genome Res. 33, 435–447 (2023).

87. Rajan-Babu, I.-S., Dolzhenko, E., Eberle, M. A. & Friedman, J. M. Sequence composition changes in short tandem repeats: heterogeneity, detection, mechanisms and clinical implications. Nat. Rev. Genet. 25, 476–499 (2024).

88. Adam, C. L., Rocha, J., Sudmant, P. & Rohlfs, R. TRACKing tandem repeats: a customizable pipeline for identification and cross-species comparison. Bioinform. Adv. 5, vbaf066 (2025).

89. Yoo, D., et al. Complete sequencing of ape genomes. bioRxivorg (2024) doi:10.1101/2024.07.31.605654.

90. Benson, G. Tandem repeats finder: a program to analyze DNA sequences. Nucleic Acids Res. 27, 573–580 (1999).

91. Hubley, R. et al. The Dfam database of repetitive DNA families. Nucleic Acids Res. 44, D81–9 (2016).

92. Rodriguez, J. M. et al. APPRIS: annotation of principal and alternative splice isoforms. Nucleic Acids Res. 41, D110–7 (2013).

93. Hinrichs, A. S. et al. The UCSC Genome Browser Database: update 2006. Nucleic Acids Res. 34, D590–8 (2006).

94. Madeira, F. et al. The EMBL-EBI Job Dispatcher sequence analysis tools framework in 2024. Nucleic Acids Res. 52, W521–W525 (2024).

95. Liao, W.-W. et al. A draft human pangenome reference. Nature 617, 312–324 (2023).

96. Dolzhenko, E. et al. Characterization and visualization of tandem repeats at genome scale. Nat. Biotechnol. 42, 1606–1614 (2024).

97. Shao, Y. et al. Phylogenomic analyses provide insights into primate evolution. Science 380, 913–924 (2023).

98. Ge, S. X., Jung, D. & Yao, R. ShinyGO: a graphical gene-set enrichment tool for animals and plants. Bioinformatics 36, 2628–2629 (2020).

99. Ritchie, M. E. et al. limma powers differential expression analyses for RNA-sequencing and microarray studies. Nucleic Acids Res. 43, e47 (2015).

