## supplementary_figures for "Tandem repeat variation within and between species reveals signatures of selection in humans and chimpanzees"

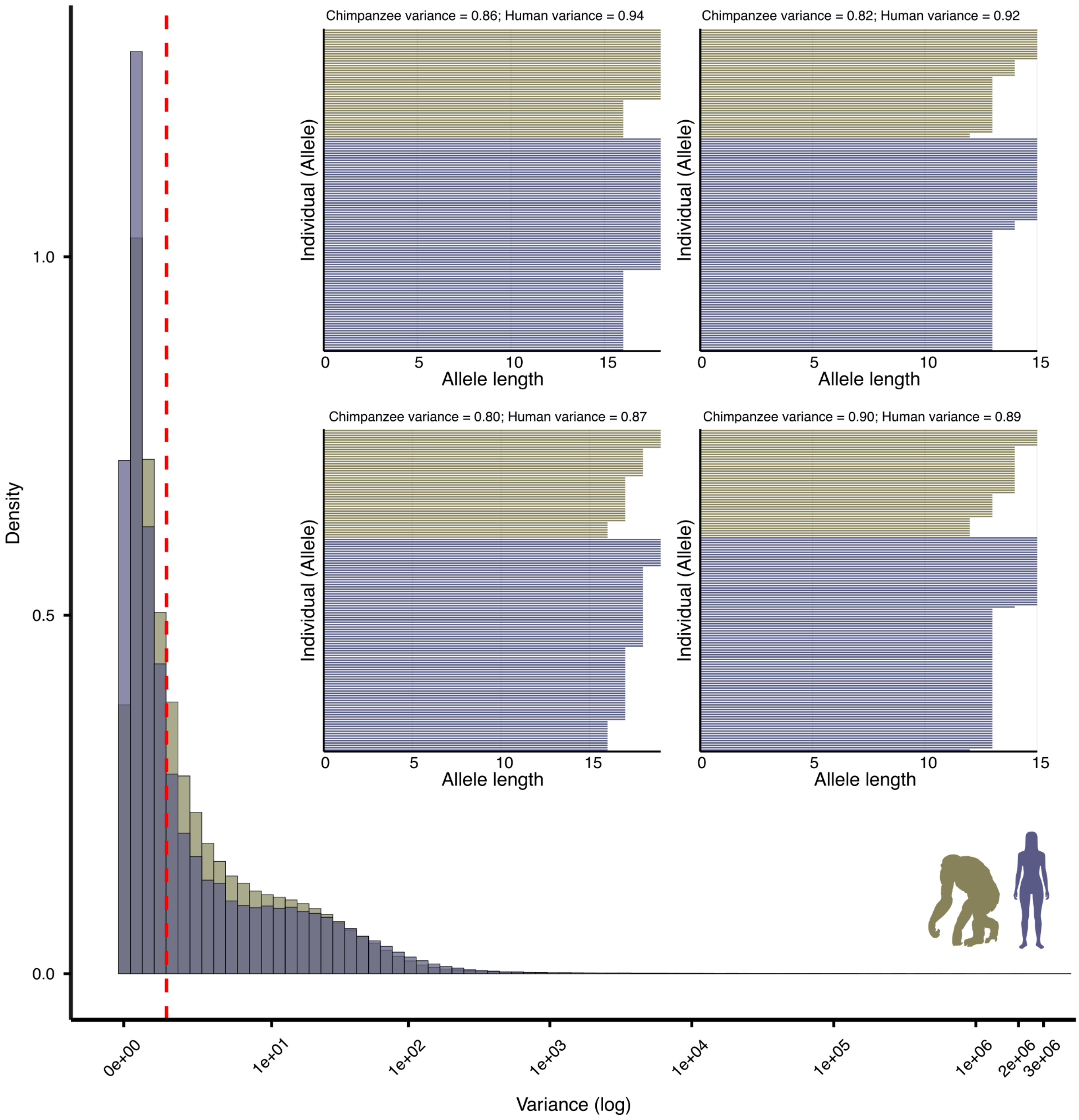


Supplementary Figure 1 - Histogram showing the distribution of non-zero within-species tandem repeat (TR) variance across humans and chimpanzees. The vertical line indicates the implemented threshold (minimum variance = 1), used to exclude loci with low variance that yield small ratios, which are likely not biologically relevant. Inset barplots show four examples of the allele-length distributions of TR variants with within-species variance near but below 1.

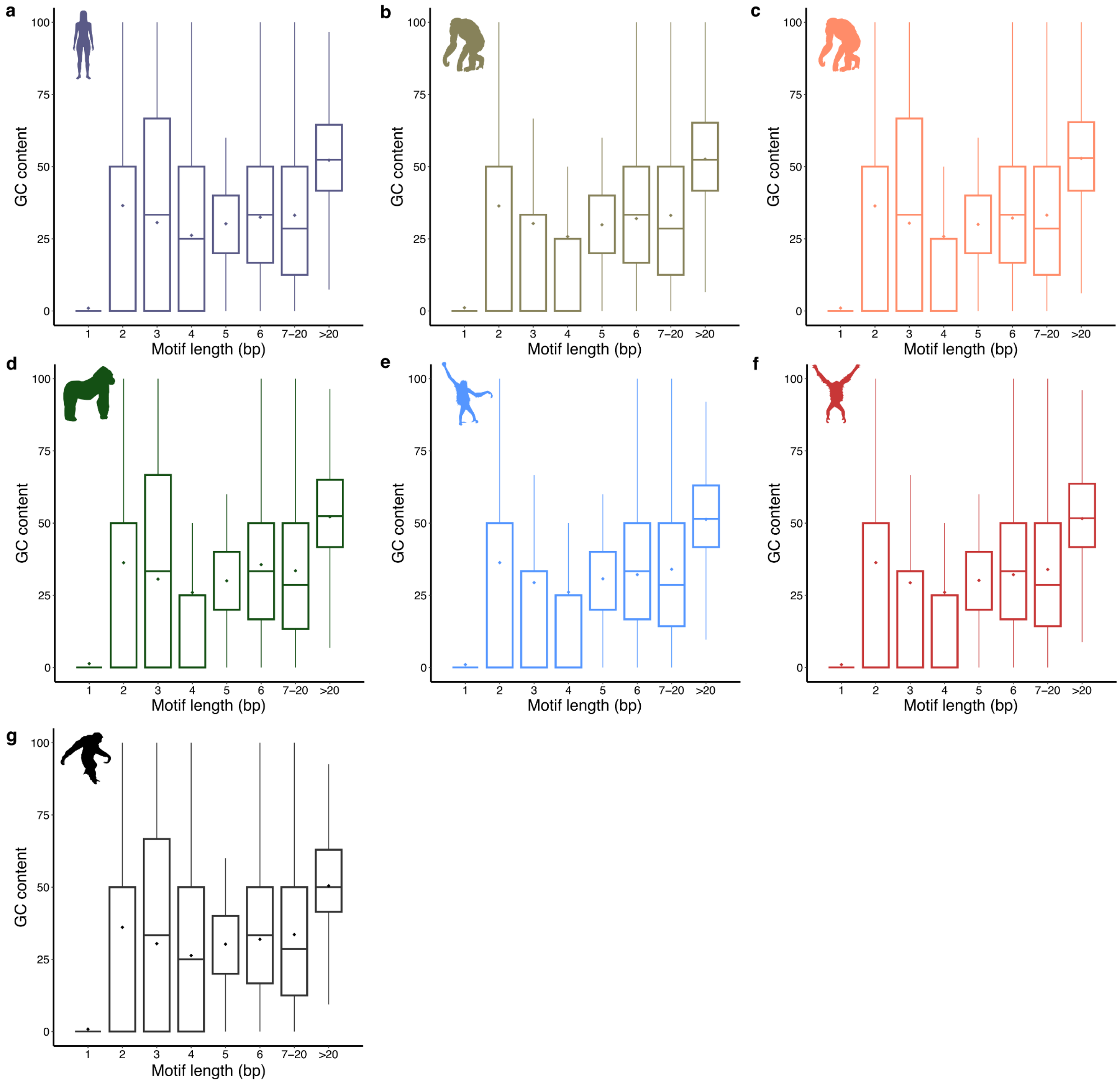


Supplementary Figure 2 - Boxplot showing the GC content of tandem repeat (TR) motifs across different motif lengths for **a**, *Homo sapiens*; **b**, *Pan troglodytes*; **c**, *Pan paniscus*; **d**, *Gorilla gorilla*; **e**, *Pongo pygmaeus*; **f**, *Pongo abelii*; **g**, *Symphalangus syndactylus;* and **h**, *Macaca mulatta.*

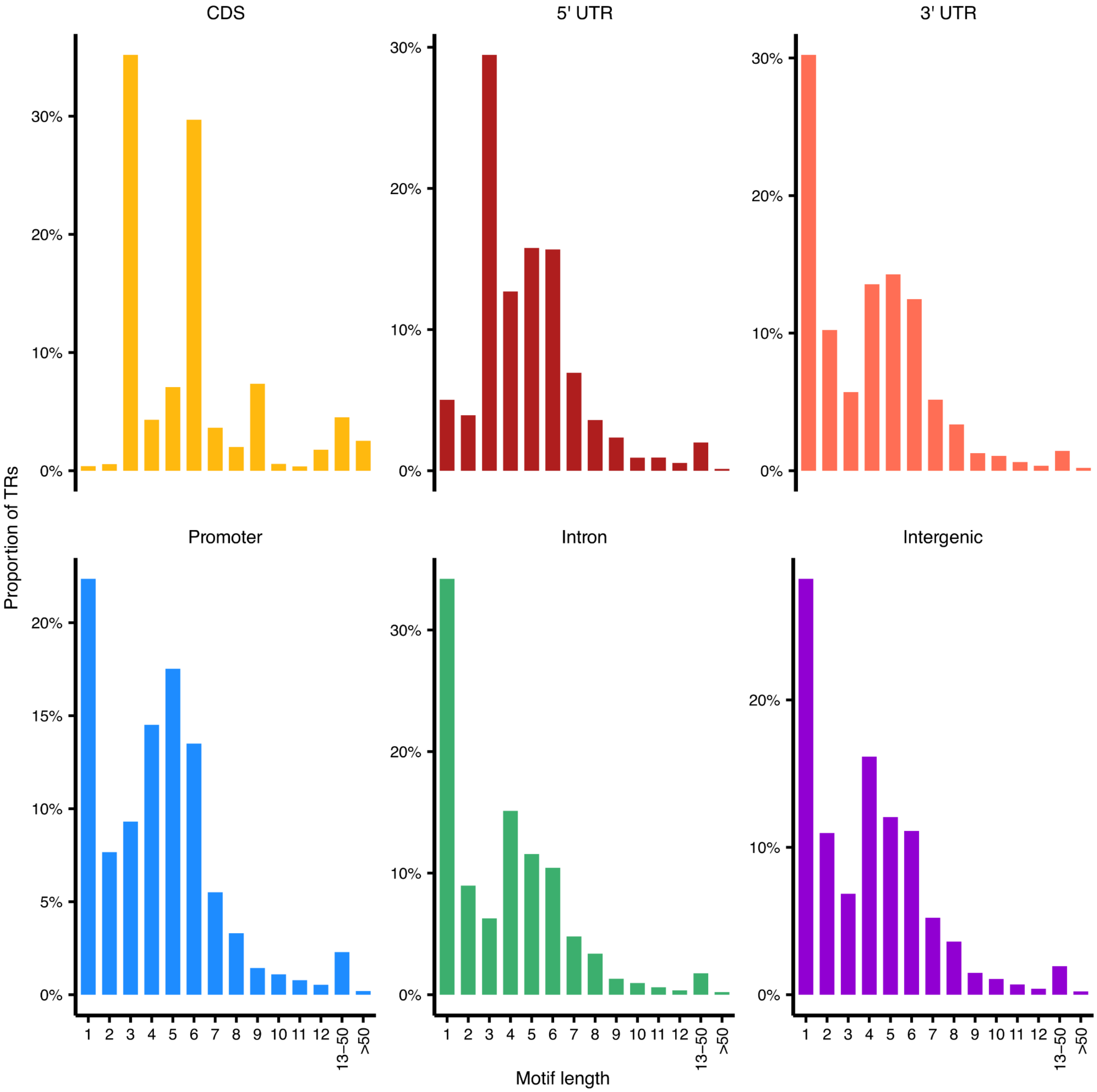


Supplementary Figure 3 - Barplots showing the proportional distribution of tandem repeat (TR) motif lengths across genomic features in the CHM13 reference genome.

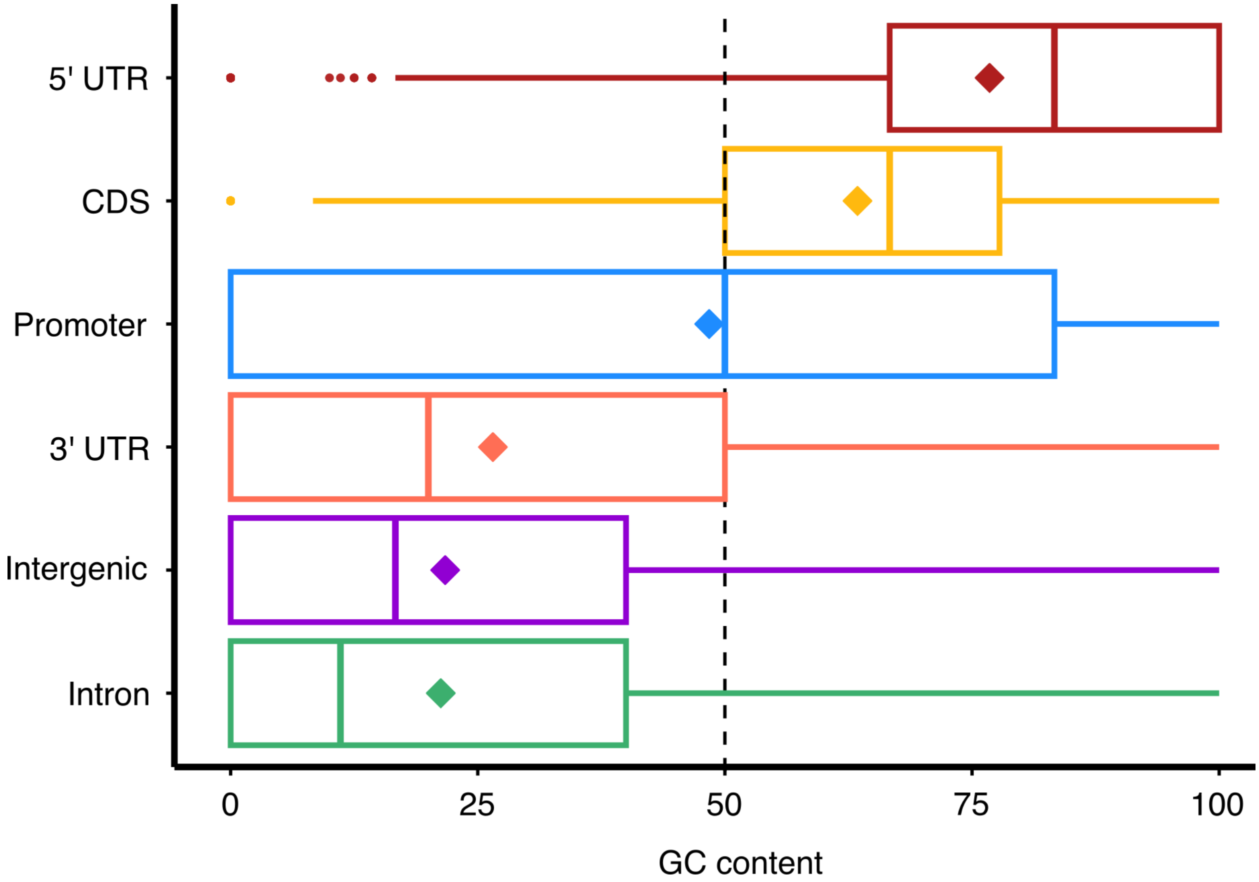


Supplementary Figure 4 - GC content of TR motifs across genomic features in the human genome.

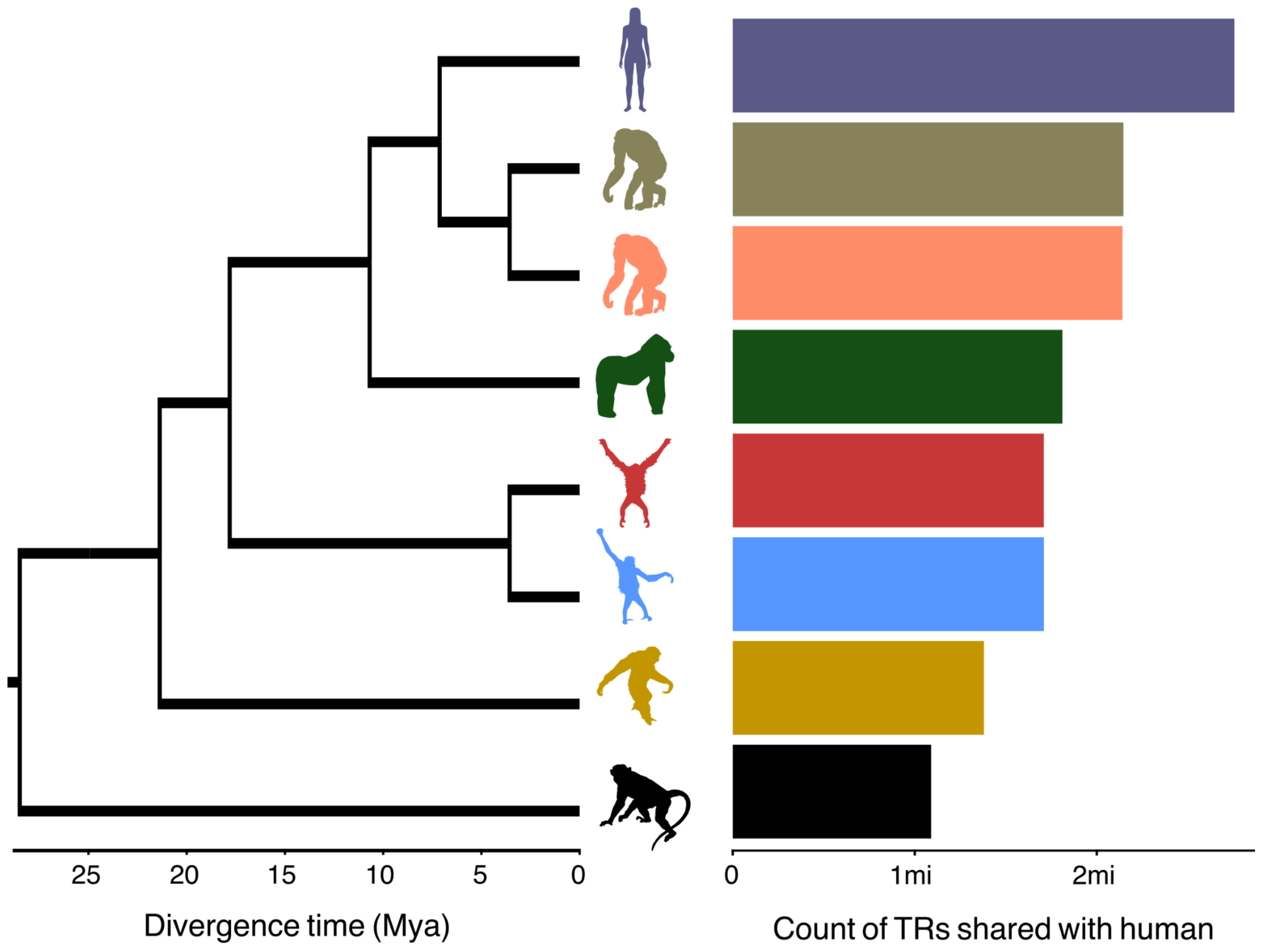


Supplementary Figure 5 - Phylogenetic tree of the seven ape genomes with a barplot showing each species’ number of homologous TRs with humans. The top bar represents the total number of TRs identified in the human genome.

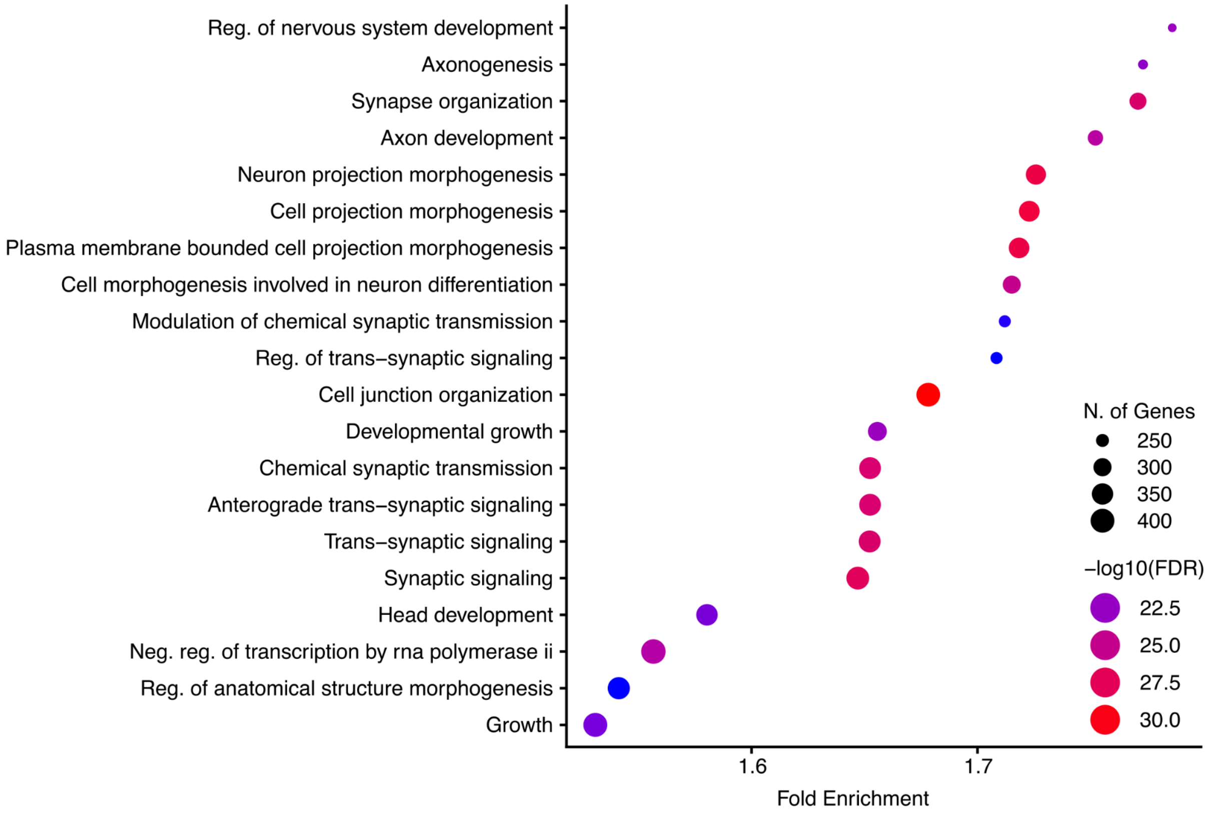


Supplementary Figure 6 - Gene Ontology enrichment analysis for biological processes terms associated with genes containing 5’ UTR TRs, using the whole set of TR-containing genes as background.

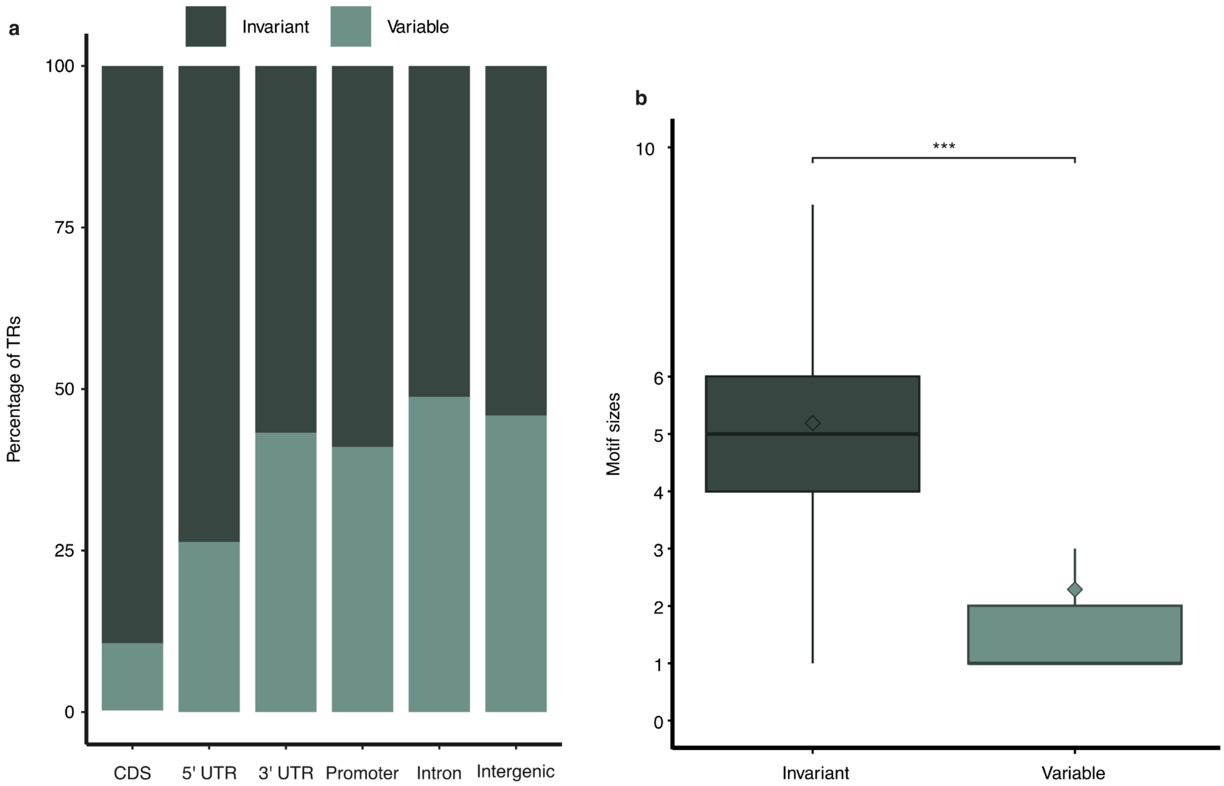


Supplementary Figure 7 - **a**, Barplots showing the percentage of invariant and variable TRs within each genomic feature. **b**, Boxplot showing the distribution of motif lengths across invariant and variable TRs. ***P ≤ 0.001 denotes statistically significant differences given by Wilcoxon rank-sum tests. Outliers identified according to Tukey's 1.5 x IQR criterion are omitted for visualisation purposes.

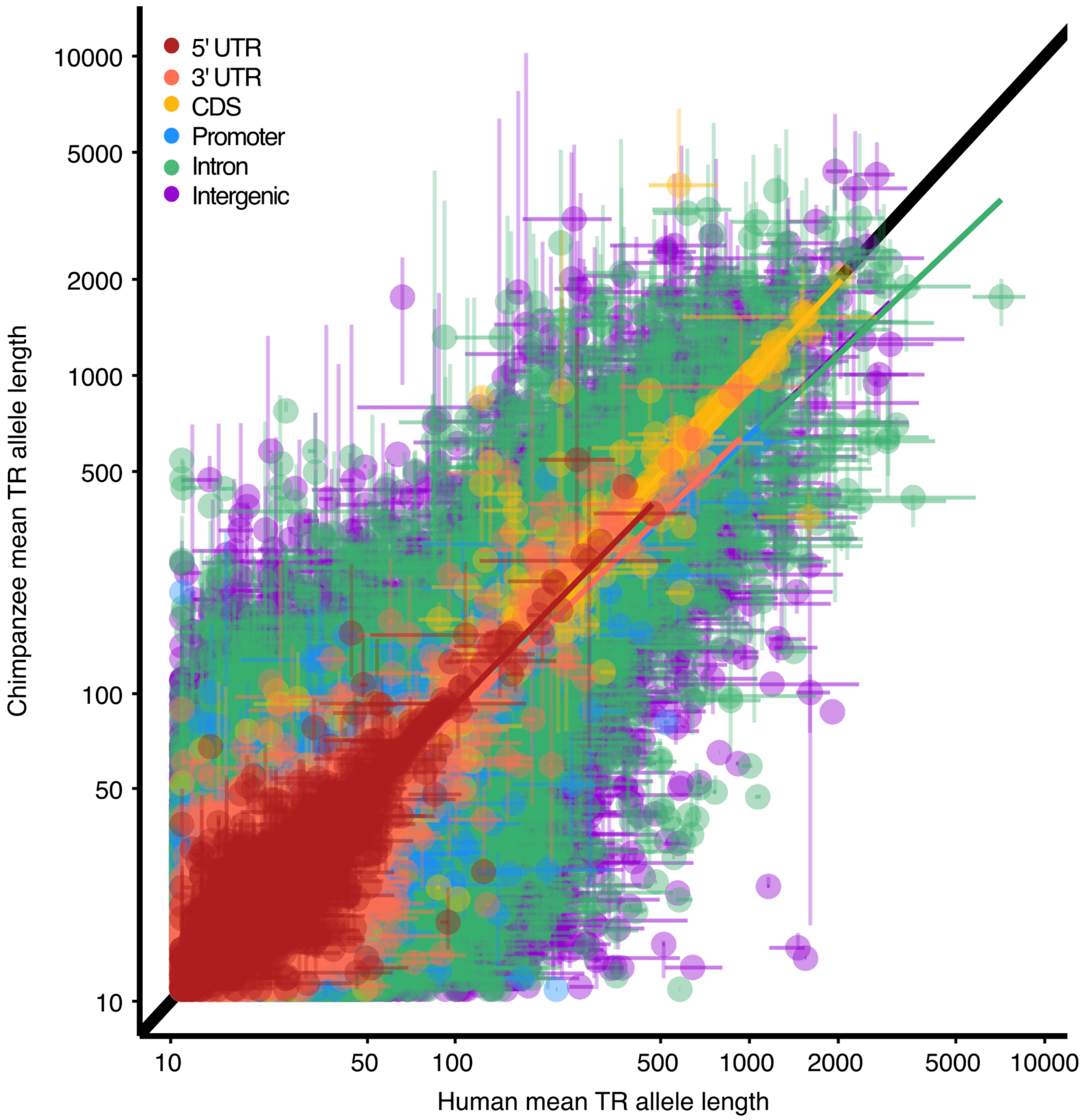


Supplementary Figure 8 - Scatterplot of mean TR allele lengths between humans (x-axis) and chimpanzees (y-axis) for TRs shared between species. Each point represents a single TR locus, color-coded by its genomic annotation in the CHM13 genome. Error bars represent the 5th-95th quantile of allele lengths in humans (horizontal bars) and chimpanzees (vertical bars).

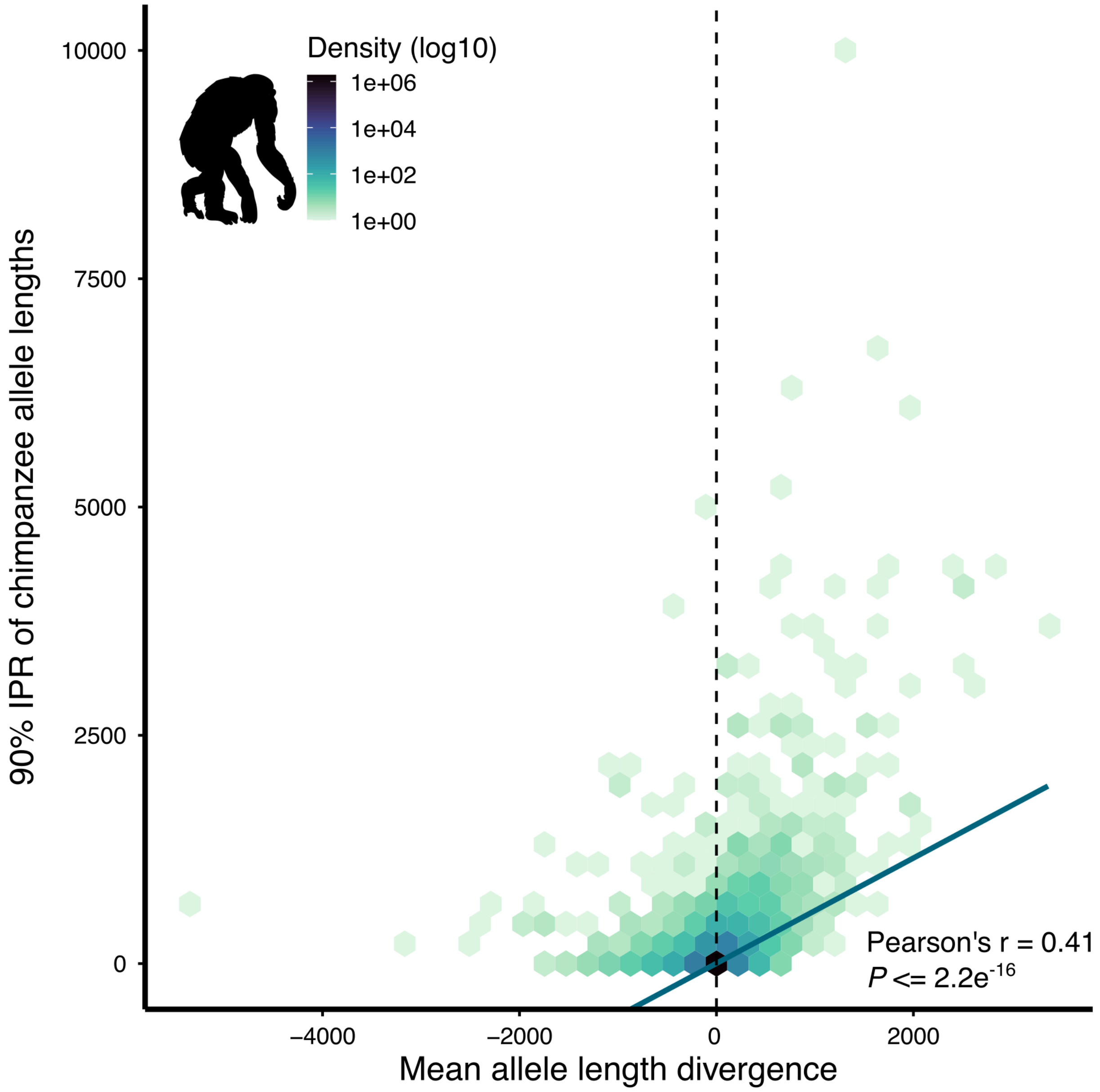


Supplementary Figure 9 - Heatmap of mean TR allele length divergence between chimpanzees and humans (y-axis) and 90% interpercentile range of chimpanzee allele lengths per locus (x-axis) across all shared TRs.

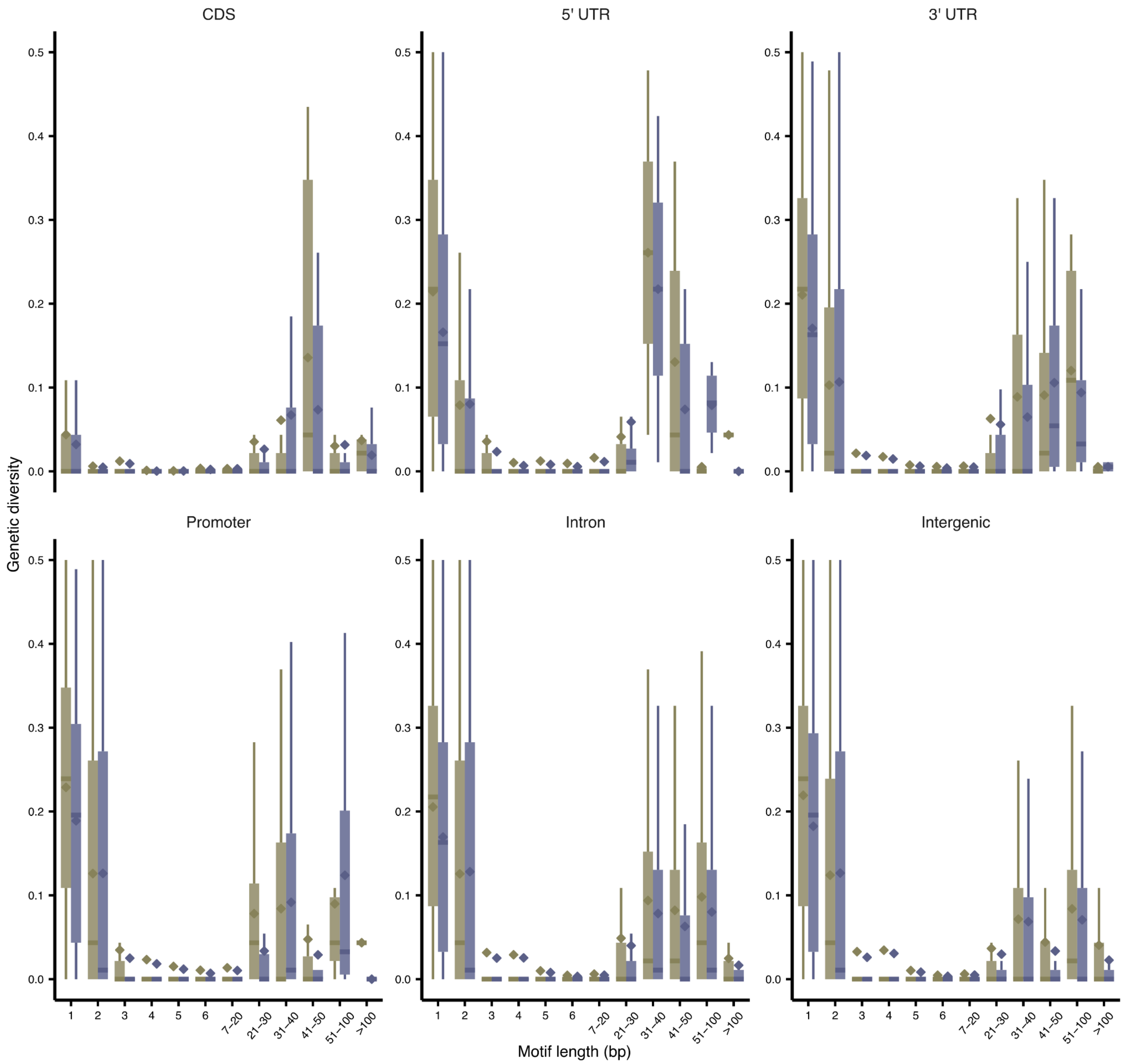


Supplementary Figure 10 - Expected heterozygosity as a proxy for genetic diversity of TRs shared between humans (purple) and chimpanzees (green) across genomic features, stratified by motif lengths.

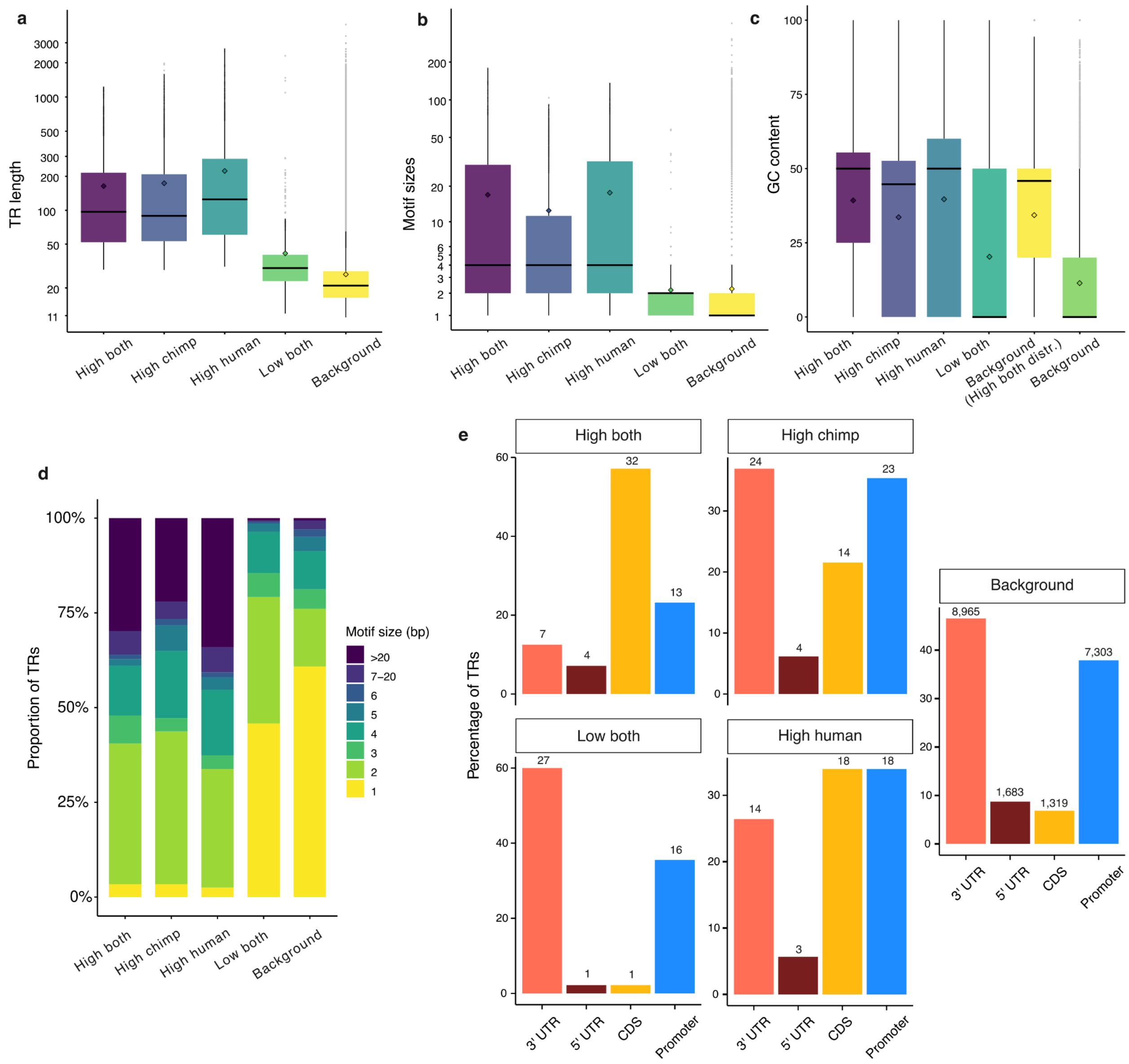


Supplementary Figure 11 - Boxplots showing the distribution of: **a**, TR lengths; **b**, motif sizes; and **c**, GC content across *D* categories. "Background HD subset" represents a subset of background TR variants (TRVs) not present in the top-ranked groups but displaying the same allele length and motif length distribution as the TRVs in the group with High D in both species. **d**, Proportions of TRs with different motif sizes across *D* categories, and **e**, proportions of genic, non-intronic TRs within each *D* category made up by different genomic features.

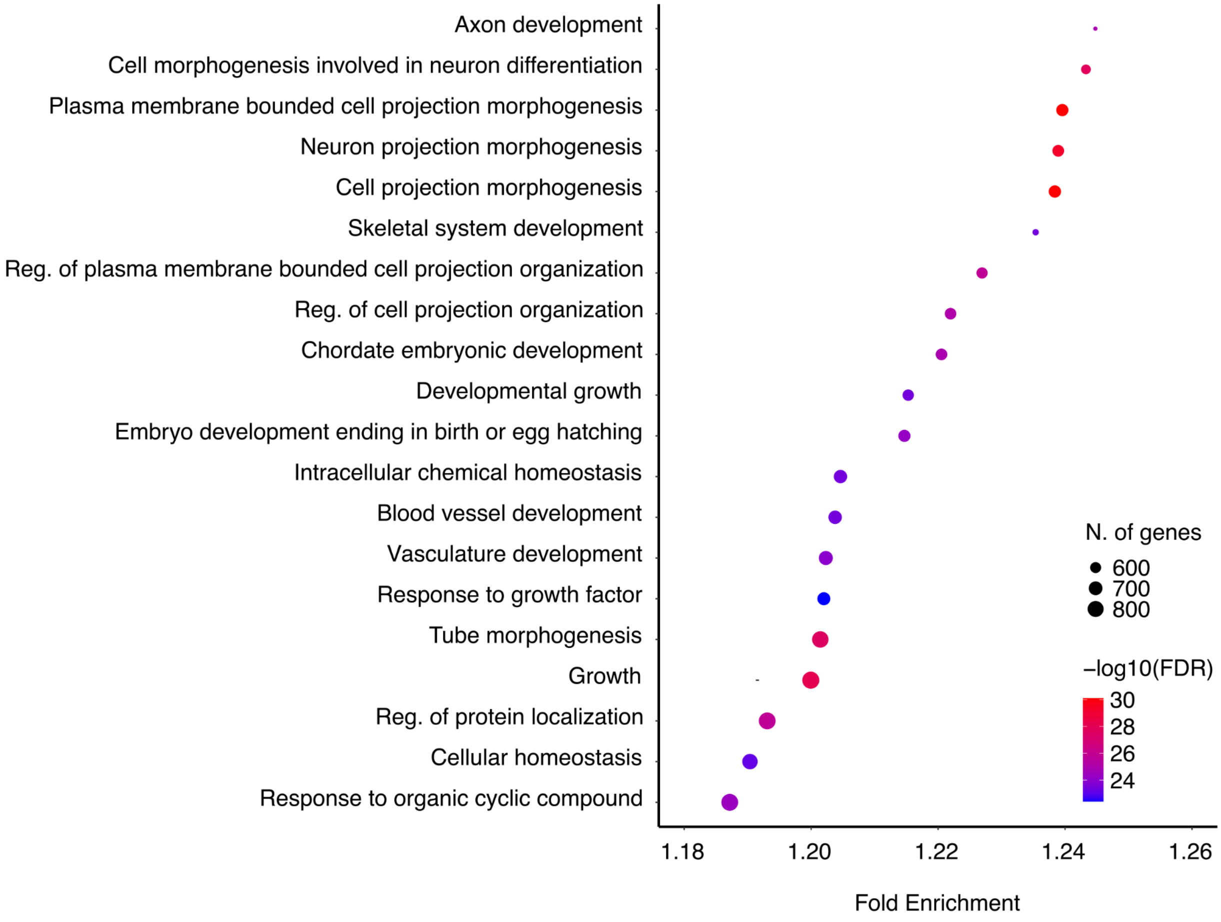


Supplementary Figure 12 - Gene Ontology enrichment analysis for biological processes terms associated with genes containing TRs using the whole set of human genes as background.

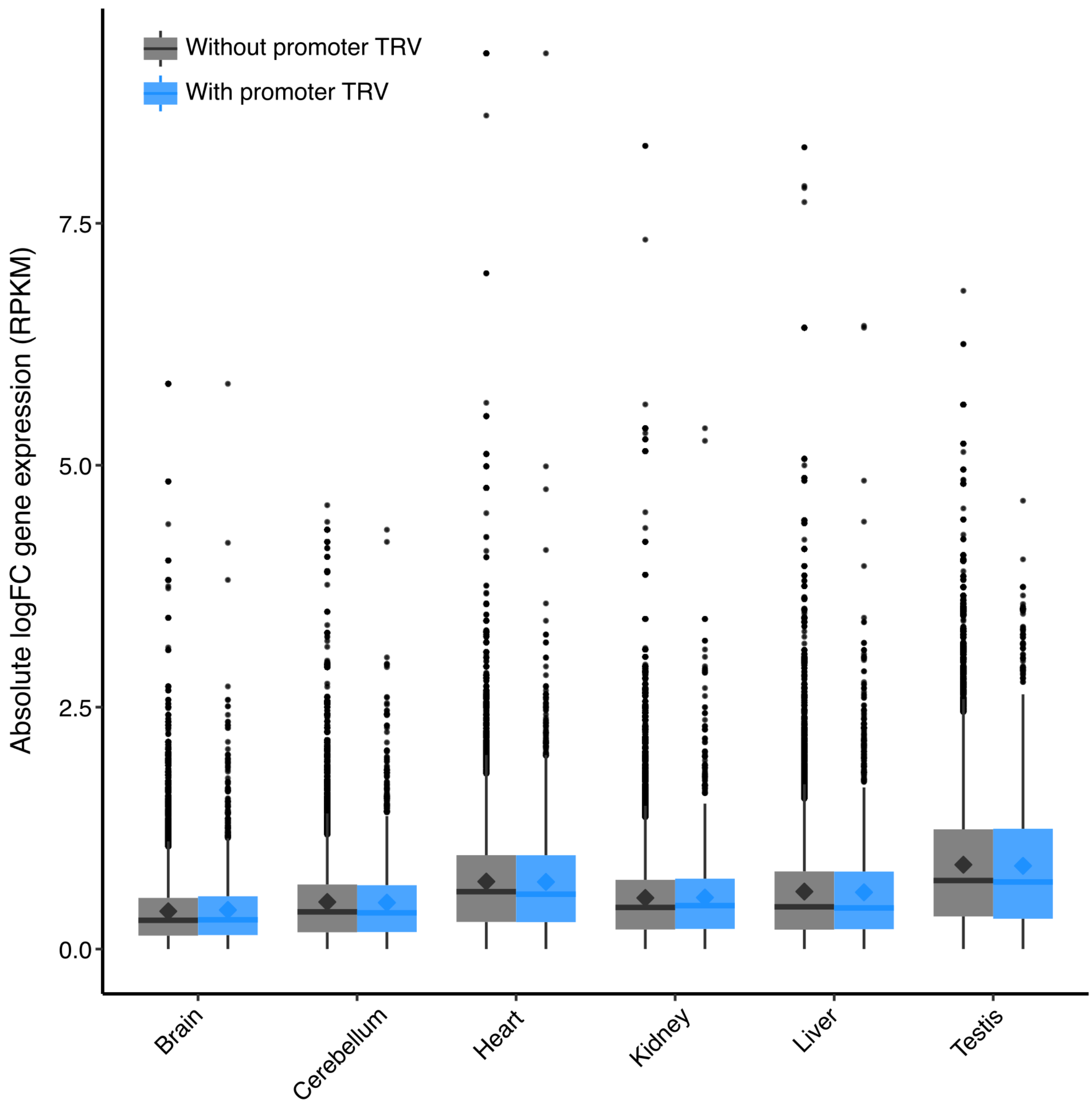


Supplementary Figure 13 - Boxplot showing the distribution of absolute log fold change in gene expression divergence between humans and chimpanzees for genes with and without promoter tandem repeat variants (TRVs) across multiple tissues.

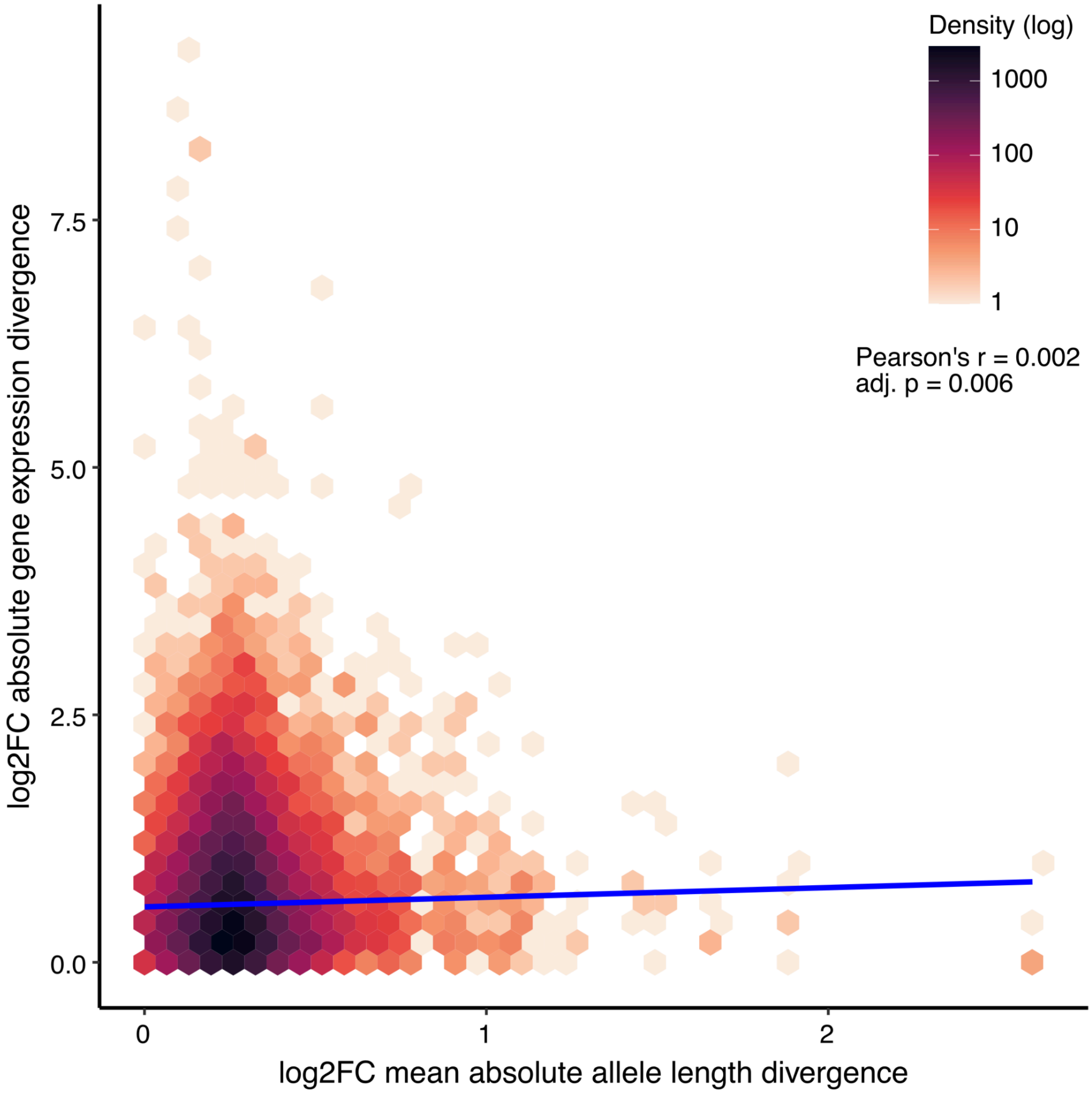


Supplementary Figure 14 - Heatmap showing the correlation between absolute log fold change in mean TR allele length averaged across genes (x-axis) and absolute log fold change in gene expression divergence between humans and chimpanzees (y-axis) across multiple tissues from Brawand et al. (2011).

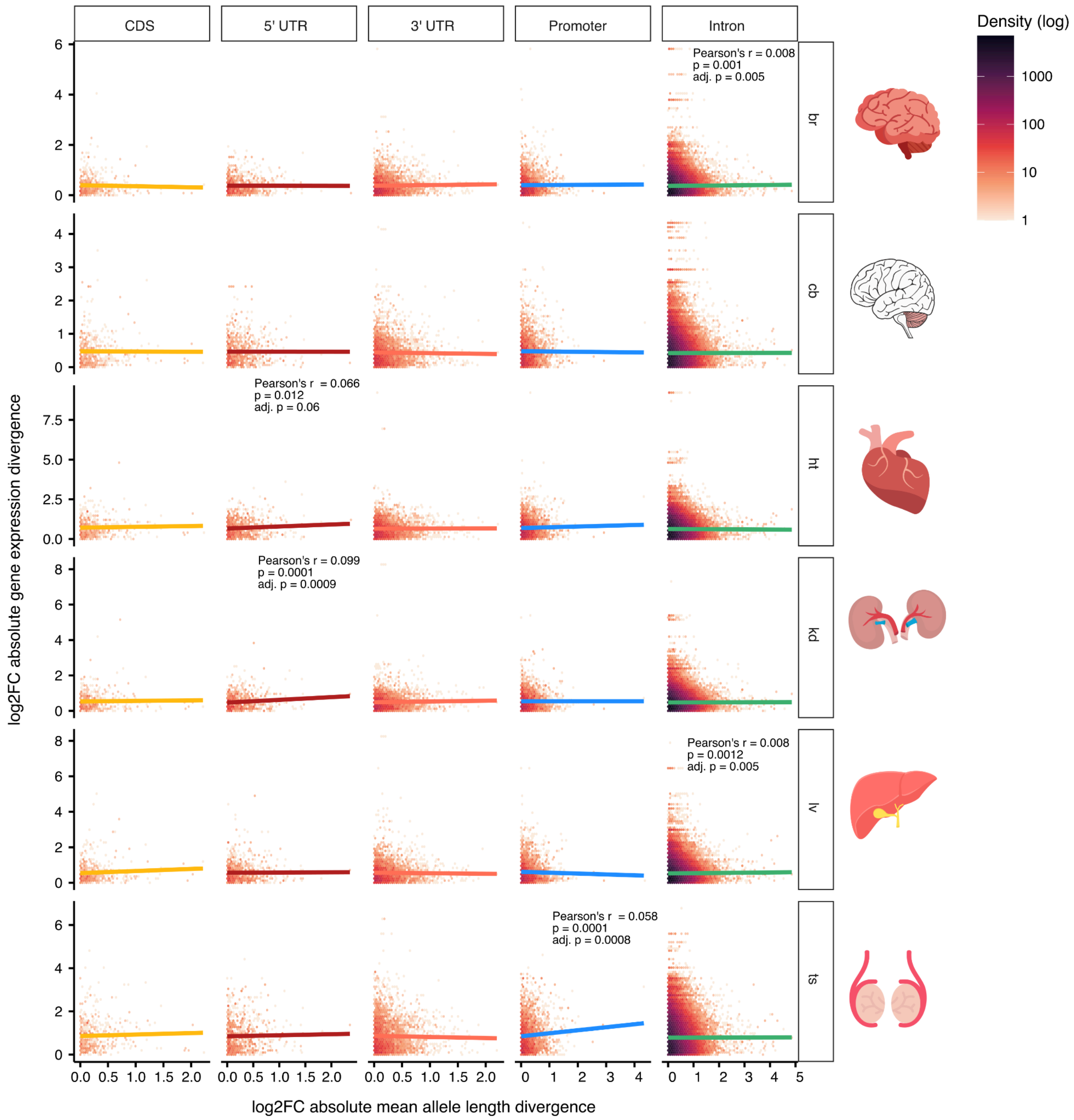


Supplementary Figure 15 - Heatmaps showing the correlation between absolute log fold change in mean tandem repeat (TR) allele length across genes (x-axis) and absolute log fold change in gene expression divergence between humans and chimpanzees (y-axis) across multiple tissues from Brawand et al. (2011).

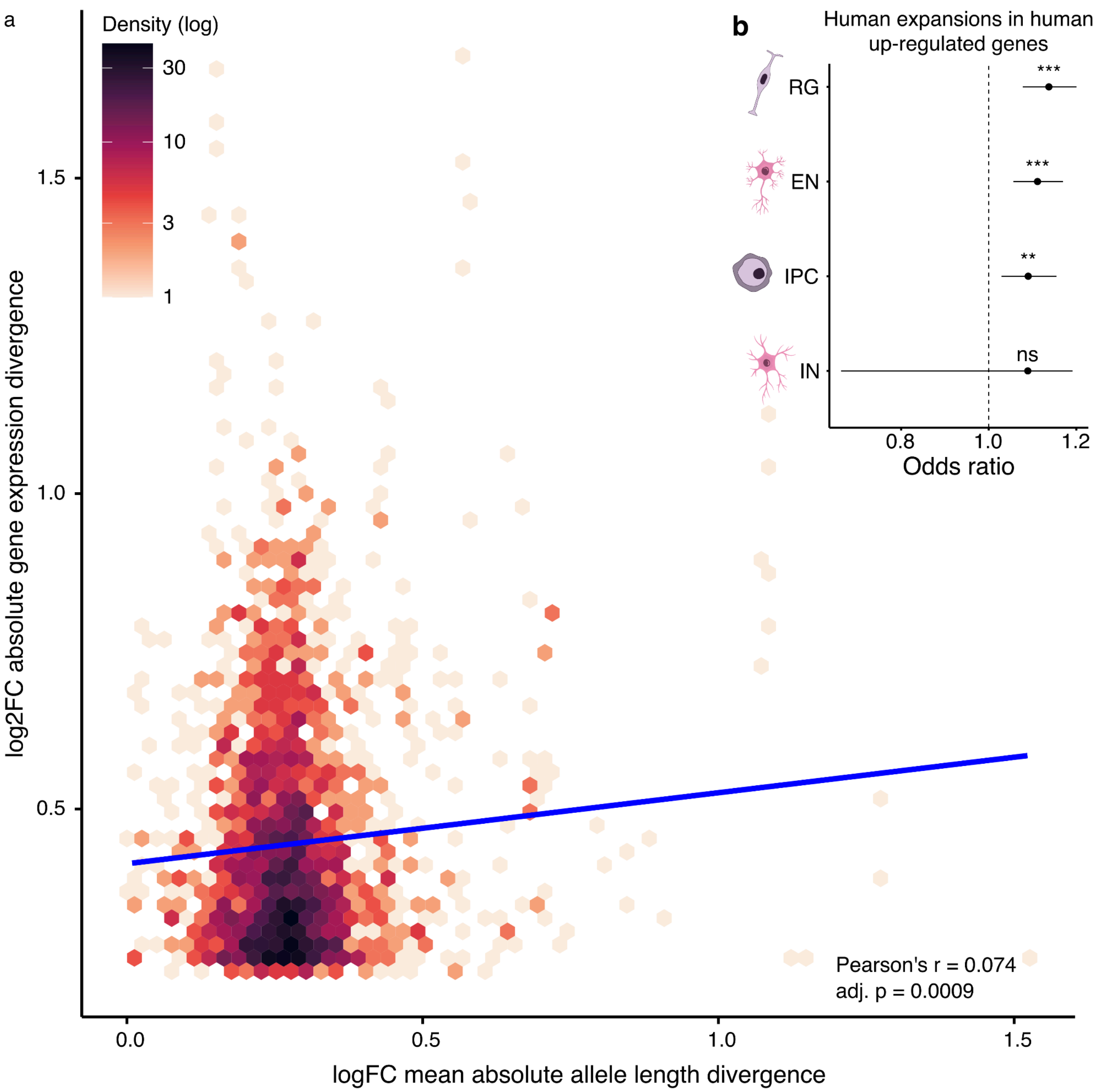


Supplementary Figure 16 - **a**, Heatmap showing the correlation between absolute log fold change in mean tandem repeat (TR) allele length averaged across genes (x-axis) and absolute log fold change in gene expression divergence between humans and chimpanzees (y-axis) across multiple organoids with telencephalon identity from Pollen et al. (2019). **b**, Odds ratio for enrichment analysis of TRs with human expansions in genes up-regulated in humans compared to genes up-regulated in chimpanzees.

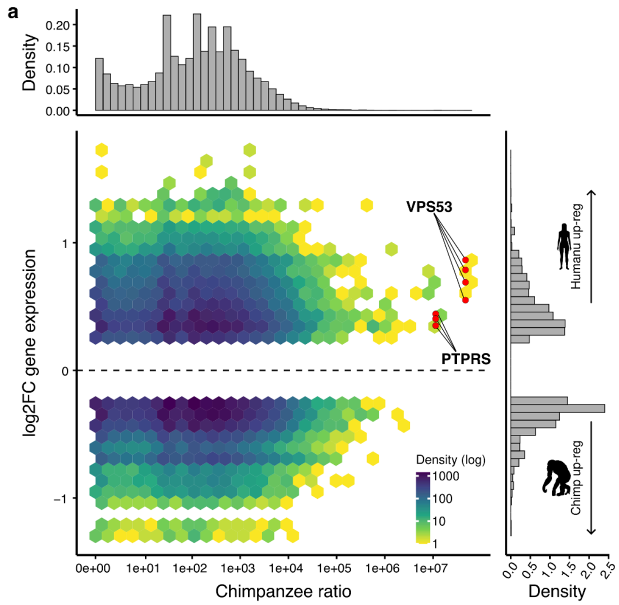


Supplementary Figure 17 - Distribution of **a**, chimpanzee and **b**, human tandem repeat variant (TRV) ratio D (x- axis) for TRs overlapping genes with different levels of expression divergence (y-axis) across cell types from primary samples or organoids with telencephalon identity (Pollen et al. 2019). Red points represent divergent TRVs that overlap genes VPS53 and PTPRS, with expression values from each of the four cell types. These genes have human-specific regulatory changes during cortical development.

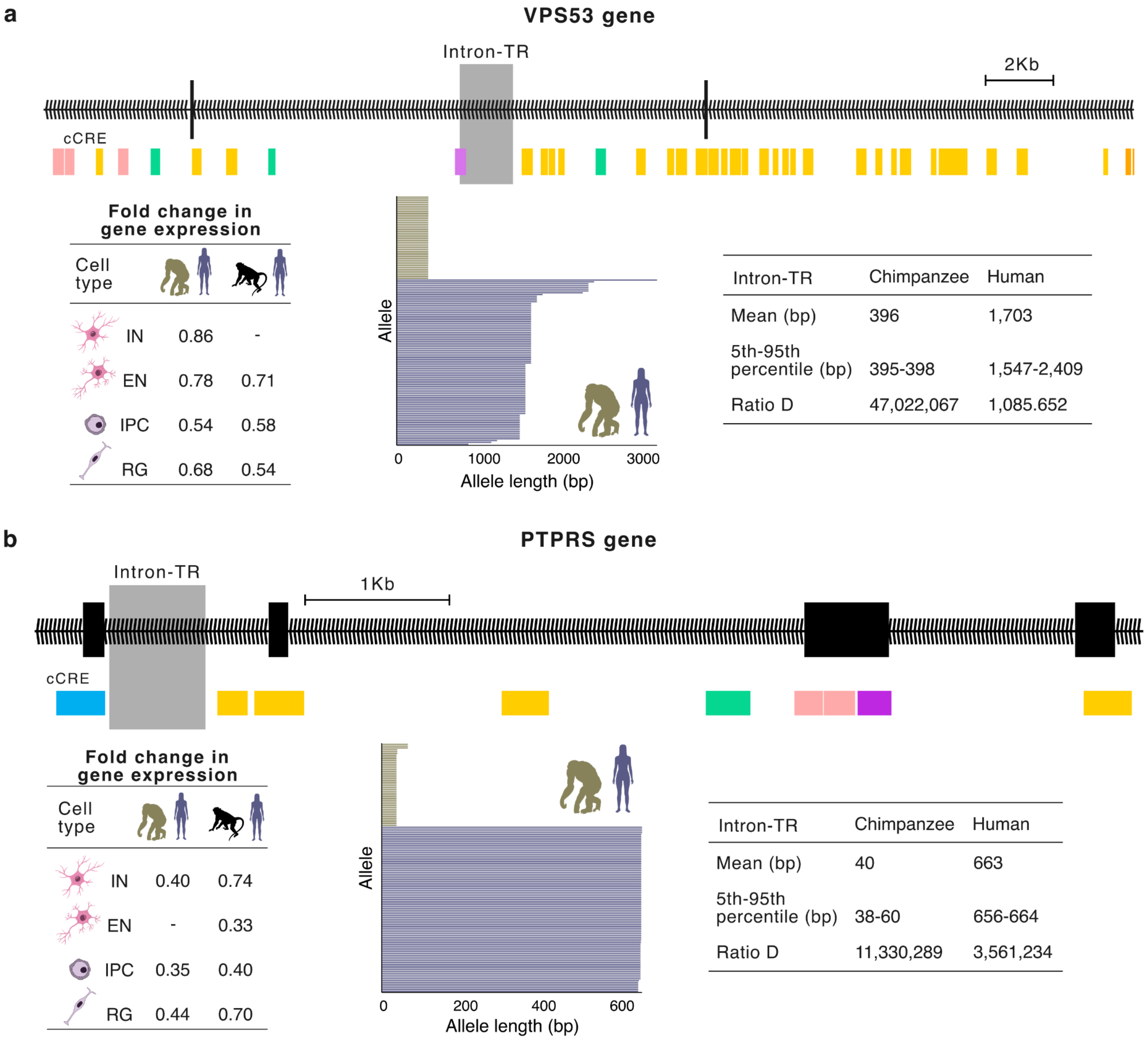


Supplementary Figure 18 - Genomic location and allele length distribution for two highly divergent intronic TRVs overlapping **a**, VPS53 and **b**, PTPRS. These genes exhibit human-specific regulatory changes during cortical development, as indicated by gene expression divergence between human and chimpanzee organoids with telencephalon identity and human and macaque cells from primary telencephalon samples. Candidate cis-regulatory elements (cCREs) indicate proximal (orange) and distal (yellow) enhancer-like signatures, chromatin accessibility (CA) + H3K4me3 (pink), CA-CTCF (blue), CA + transcription factor (dark purple), CA only (green), and transcription factor only (light purple).

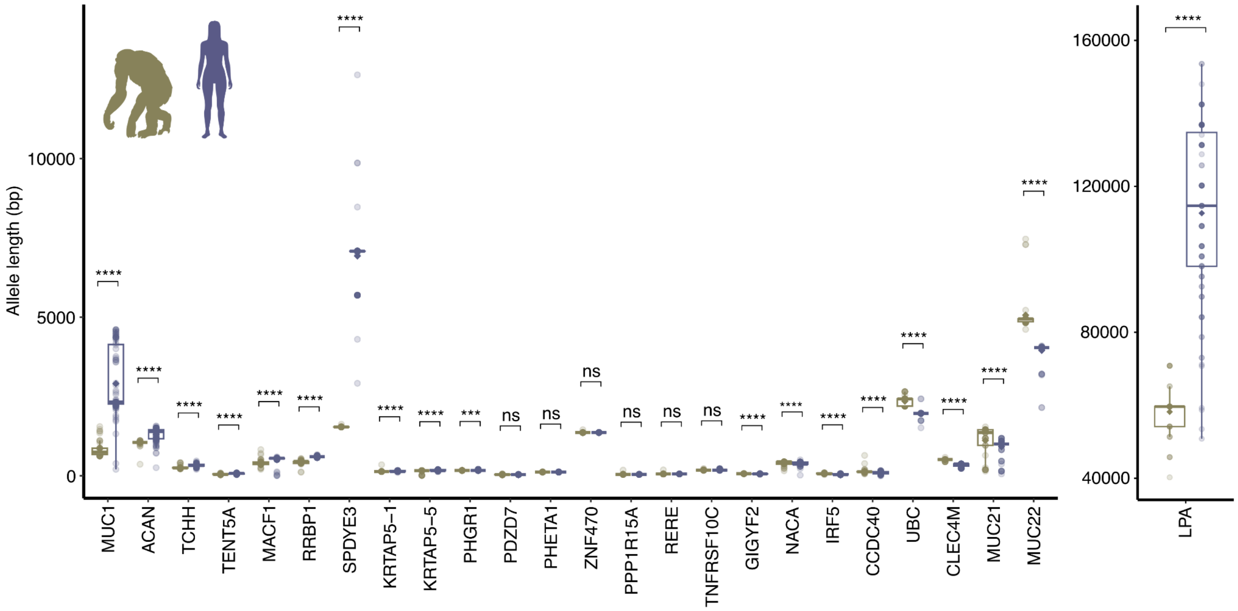


Supplementary Figure 19 - Boxplots showing the distribution of chimpanzee and human allele lengths for genic VNTRs potentially associated with phenotypes listed in the UK Biobank. *$p \leq 0.05$, **$p \leq 0.01$, ***$p \leq 0.001$, ****$p \leq 0.0001$ denote significant differences given by the Wilcoxon rank-sum test.

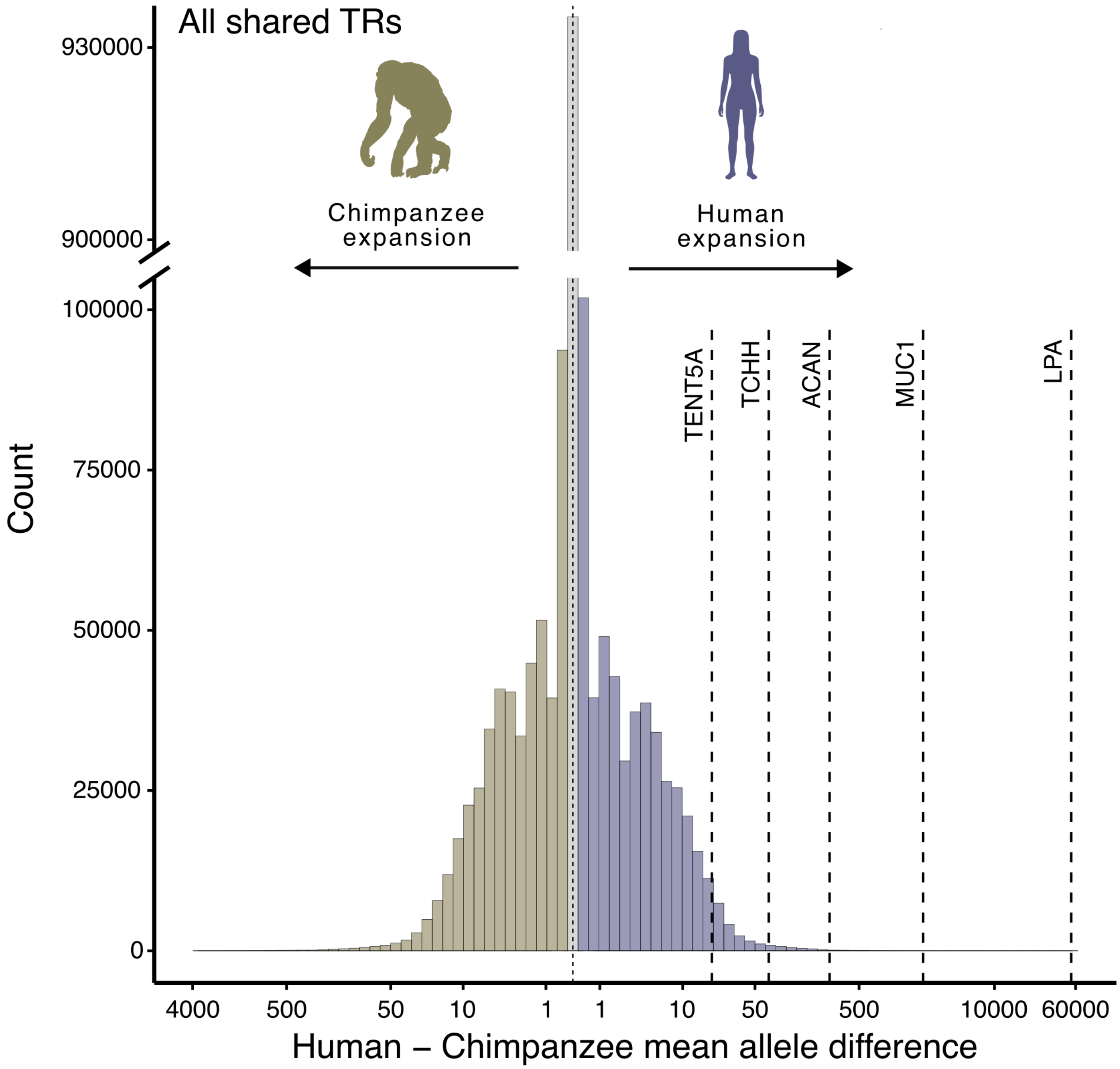


Supplementary Figure 20 - Barplot showing the distribution of mean allele length differences between humans and chimpanzees for all shared tandem repeats (TRs), highlighting five genes with trait-associated TRs in humans. Mean allele length difference between humans and chimpanzees for trait-associated TRs was significantly greater than expected under the null distribution for all TRs (permutation test, 1,000,000 runs, $p \leq 1 \times 10^{-6}$, and $p = 4.4 \times 10^{-5}$ when excluding the long and highly divergent VNTR in LPA).
